# Dissociable roles for Anterior Cingulate Cortex and Basolateral Amygdala in Decision Confidence and Learning under Uncertainty

**DOI:** 10.1101/655860

**Authors:** A Stolyarova, M Rakhshan, Evan E. Hart, Thomas J. O’Dell, MAK Peters, H Lau, A Soltani, A Izquierdo

## Abstract

It has been suggested the subjective sense of certainty, or confidence, in ambiguous sensory cues can alter the interpretation of reward feedback and facilitate learning. We trained rats to report the orientation of ambiguous visual stimuli according to a spatial stimulus-response rule. Following choice, rats could wait a self-timed delay for reward or initiate a new trial. Waiting times increased with discrimination accuracy, demonstrating that this measure could be used as a proxy for confidence. Chemogenetic silencing of BLA shortened waiting times overall whereas ACC inhibition rendered waiting times insensitive to confidence-modulating attributes of visual stimuli, suggesting contribution of ACC but not BLA to confidence computations. Subsequent reversal learning was enhanced by confidence. Both ACC and BLA inhibition blocked this enhancement but via differential modulation of learning strategies and consistency in using learned rules. Altogether, we demonstrate dissociable roles for ACC and BLA in transmitting confidence and learning under uncertainty.

## Introduction

Learning relies on the ability to use external cues to predict the state of the world, take actions based on those predictions, and associate those actions with subsequent reward. Learning such associations can be straightforward when stimuli that precede actions or rewards can be discriminated clearly. However, this is not the case in naturalistic settings in which sensory cues or stimuli are ambiguous and thus the perception of or prediction about the state of the world is uncertain. In such situations, stimulus detection or discrimination are frequently accompanied by a sense of certainty, or confidence, in choice (Grimaldi, Lau, & Basso, 2015; Kepecs, Uchida, Zariwala, & Mainen, 2008). Recent evidence indicates that confidence may influence neural activity in brain regions involved in orchestrating reward responses (Guggenmos, Wilbertz, Hebart, & Sterzer, 2016; Hebart, Schriever, Donner, & Haynes, 2016) particularly when reward is significantly delayed (Iigaya, 2016). Consequently, the sensory properties of reward-predicting cues and confidence in disambiguating them may directly influence valuation (Chen, Mihalas, Niebur, & Stuphorn, 2013) and learning from reward feedback.

Recent studies in humans have revealed neural correlates of confidence estimation and learning in several brain regions, including the prefrontal cortex (Morales, Lau, & Fleming, 2018; Rounis, Maniscalco, Rothwell, Passingham, & Lau, 2010). However, it is unclear whether these areas are causally involved in these processes. Despite powerful interference techniques in rodents (Mahler & Aston-Jones, 2018; Smith, Bucci, Luikart, & Mahler, 2016), most rodents studies on neural mechanisms of confidence have been conducted within olfactory and auditory modalities (Foote & Crystal, 2007; Lak et al., 2014). In contrast, human studies on choice and learning under perceptual uncertainty have focused on visual processing, making it difficult to link findings across species.

Here, we trained rats to report the orientation of noisy Gabor patches by making spatial choices based on a learned stimulus-response rule (e.g., Horizontal→left and Vertical→right). We manipulated different aspects of the visual stimuli to alter performance and uncertainty associated with discriminating the orientation. Following action selection using a touchscreen, rats expressed their confidence by time-wagering: they could wait for a variable amount of time before they could receive a possible reward or initiate a new trial (Lak et al., 2014). This design allowed us to measure confidence on a trial-by-trial basis. After ensuring that rats learned the stimulus-response associations, we reversed these associations to study the effect of confidence on learning using a counterbalanced design. Extensive studies in rodents have shown a distributed network supports learning and choice involving uncertain outcomes, including basolateral amygdala, BLA (Ghods-Sharifi, St. Onge, & Floresco, 2009; St Onge, Stopper, Zahm, & Floresco, 2012; Stopper & Floresco, 2011; Winstanley & Floresco, 2016), and anterior cingulate cortex, ACC (Akam et al., 2017; Tervo et al., 2014). Therefore, we used inhibitory Designer Receptors Exclusively Activated by Designer Drugs (DREADDs) to transiently inactivate projection neurons in the ACC or BLA in order to test the causal role of each area in confidence estimation or computation, and learning under perceptual uncertainty.

## Results

### Waiting time provides a proxy for confidence

To assess confidence during perceptual choice with uncertain visual information, we used a novel experimental paradigm in which rats were first presented with a single Gabor patch with one of two possible dominant orientations (horizontal (H) and vertical (V)) embedded in noise (**Figure 1A, B**). Perceptual uncertainty was manipulated using two parameters of the visual stimuli: 1) the signal-to-noise ratio (SNR), defined as the ratio of the contrast of the Gabor patch relative to the contrast of the added Gaussian noise; and 2) the overall contrast of *both* the Gabor patch *and* the added noise for a given SNR. These manipulations allowed us to modulate performance and confidence independently in order to design “matched-performance different-confidence” stimulus pairs used for learning (Grimaldi, Cho, Lau, & Basso, 2018; Koizumi, Maniscalco, & Lau, 2015; Odegaard et al., 2018). Following stimulus presentation, rats reported the perceived orientation by nosepoking one of the two side compartments of the touchscreen based on a complementary stimulus-response rule (e.g., H→left and V→right). Following action selection, rats expressed confidence in their response via time wagering; that is, they could wait for a probabilistically delivered reward if confident or initiate a new trial otherwise (see Methods).

**Figure 1.**
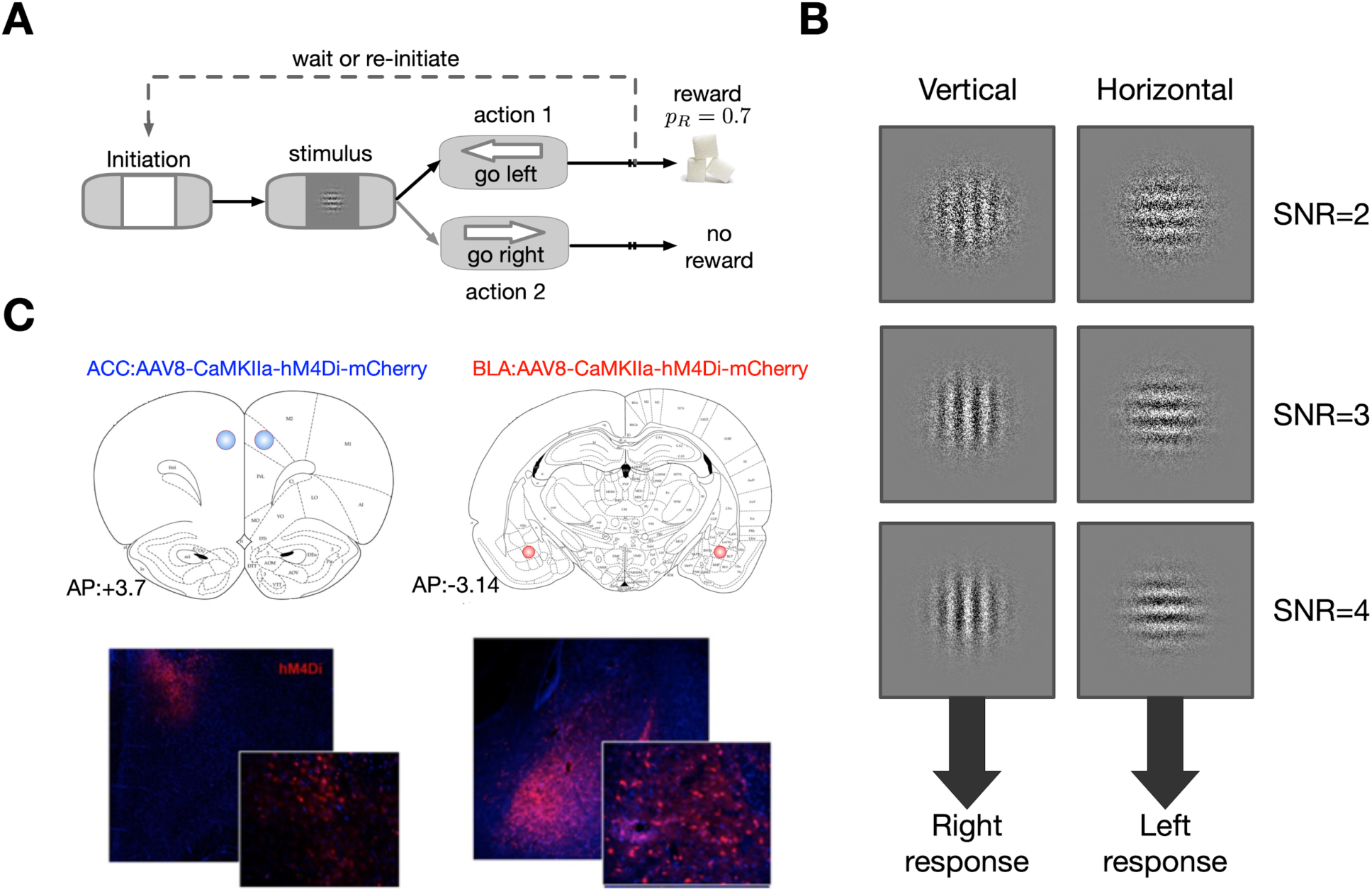
Schematic of the experimental paradigm, visual stimuli, stimulus-response rule, and DREADDs expression. **(A)** Timeline of each trial in the learning under perceptual uncertainty task. The rat first initiated a trial by nosepoking a white square in the center of the screen. The initiation stimulus then disappeared, and the rat was briefly presented (1*s*) with a single horizontal (H) or vertical (V) Gabor patch masked by noise. Rats were required to report the dominant orientation (H or V) via nosepoke based on a complementary stimulus-response rule; e.g., H→left and V→right. Correct choices were rewarded probabilistically (on 70% of randomly selected trials), following variable delay times. After stimulus discrimination, rats could wait a self-timed delay in anticipation of reward or initiate a new trial. The initiation stimulus appeared on the touchscreen 2 seconds after a rat indicated its choice. **(B)** Examples of visual stimuli and one of the two stimulus-response rules. We refer to their discriminability as an SNR value reflecting the strength of visual signal (4, most discriminable; 3, moderately discriminable; 2, least discriminable). After discrimination of the visual stimulus, the rat makes a response (using the touchscreen) according to the rule H→Left and V→right. **(C)** Representative expression of inhibitory (Gi-coupled) DREADDs under CaMKIIa are shown in ACC and BLA.

We found that during sessions with vehicle administration, our main control condition, waiting time and discrimination performance (the signal detection theoretic metric *d*′, a reliable measure of perceptual discrimination capacity; see Methods) were modulated by both the SNR and contrast of visual stimuli (**Supplementary Figure 1**). More specifically, waiting time monotonically increased with SNR for any value of contrast (GLM; contrast = 40: *F*(7,868) = 47.9, *p* = 1.5 × 10^−57^, adjusted *R*^2^ = 0.273; SNR: *β* = 1.13, *p* = 0.0002; contrast = 60: *F*(7,864) = 94, *p* = 8.7 × 10^−102^, adjusted *R*^2^ = 0.428; SNR: *β* = 2.37, *p* = 7.7 × 10^−11^; contrast = 80: *F*(7,869) = 132, *p* = 7.3 × 10^−132^, adjusted *R*^2^ = 0.511; SNR: *β* = 2.52, *p* = 2.7 × 10^−10^) but this effect was modulated by overall stimulus contrast (GLM; *F*(15,2609) = 208, *p* = 1 × 10^−130^, adjusted *R*^2^ = 0.542; SNR × contrast: *β* = 0.03, *p* = 0.0008). Therefore, we focused our analyses on SNR by averaging over all contrast levels in order to more easily examine and illustrate the effects of perceptual uncertainty on confidence for a given performance level. Nonetheless, all our results also hold when we perform our analyses for each contrast level separately (**Supplementary Figures 2 and 3**).

**Figure 2.**
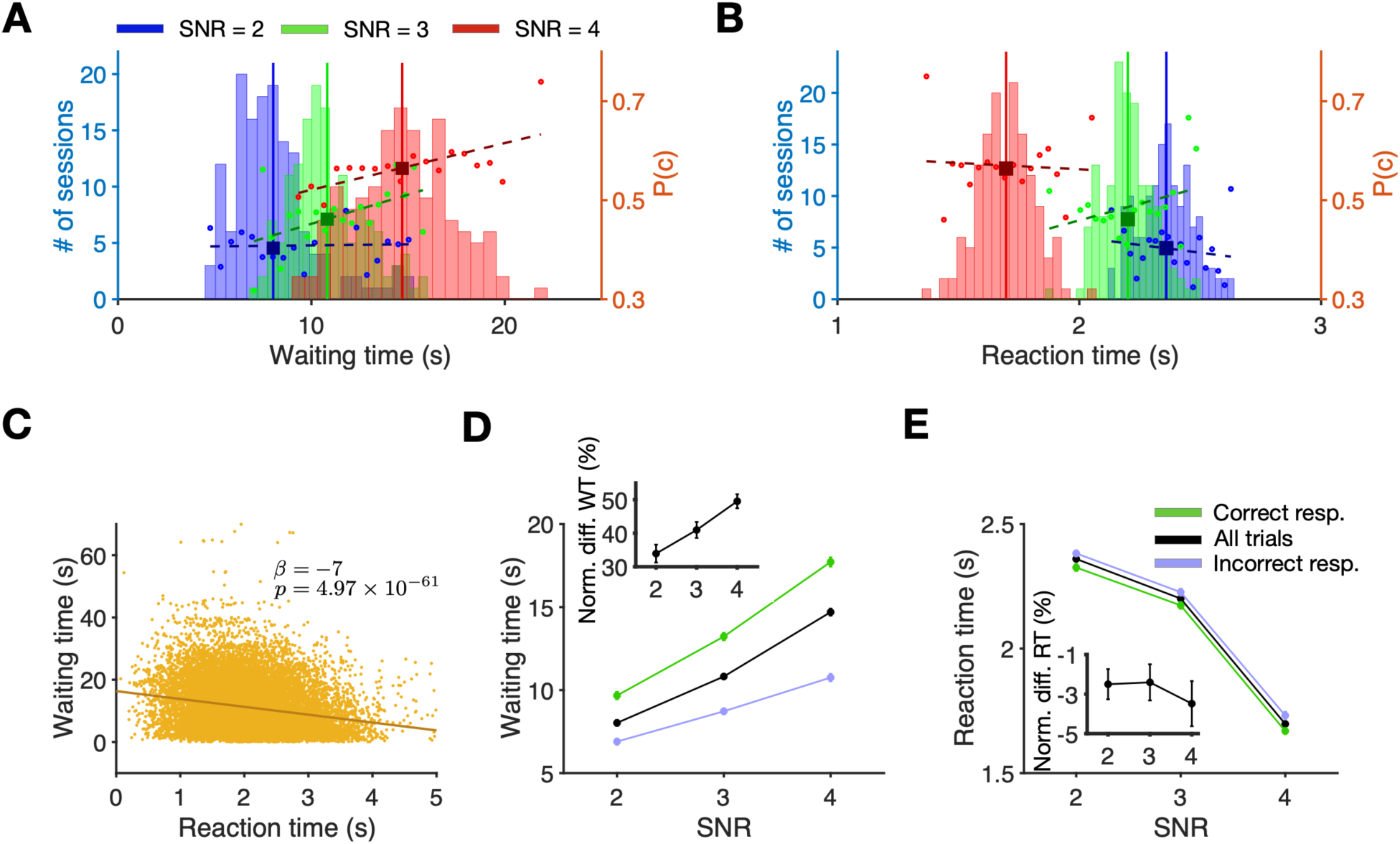
Waiting time serves as a proxy for confidence that is more sensitive than reaction time. **(A)** Waiting time before re-initiation increases as SNR increases. Plotted are the distributions of waiting times for each SNR: 2 (blue), 3 (green), and 4 (red), following vehicle administration. Solid lines show the median of each distribution. The y-axis on the right shows the probability of a correct response. Solid circles indicate the calculated probability of correct responses for each bin of the waiting time distribution, for each SNR. The dashed lines show the regression line of the probability of correct responses for bins in each SNR. **(B)** Reaction time decreases as SNR increases. Plotted are the distributions of reaction time for all trials. Similar to panel A, y-axis on the right shows the probability of a correct response. Solid circles indicate the calculated probability of correct responses for each bin of the reaction time distribution, for each SNR. Conventions are the same as in panel A. **(C)** Waiting time before re-initiation of a new trial is negatively correlated with reaction time to make a choice. Waiting time is plotted as a function of the reaction time for all trials and all rats. Each data point is a trial in a session following vehicle administration. **(D)** Waiting time is larger for correct compared to incorrect responses for any SNR. Plotted is waiting time for all trials (black), correct trials (green), and incorrect trials (purple) for different SNR. The inset shows the relative difference in waiting time between correct and incorrect responses for different SNR. Error bars show the S.E.M. over sessions (typically smaller than the symbol). **(E)** Reaction time only weakly reflects the accuracy of the response. Plotted is the reaction time for all trials and separately for correct and incorrect responses. Conventions are the same as in panel D. The inset shows the relative difference in reaction time between correct and incorrect responses for different SNR. Overall, response accuracy is reflected in waiting time an order of magnitude better than in reaction time.

**Figure 3.**
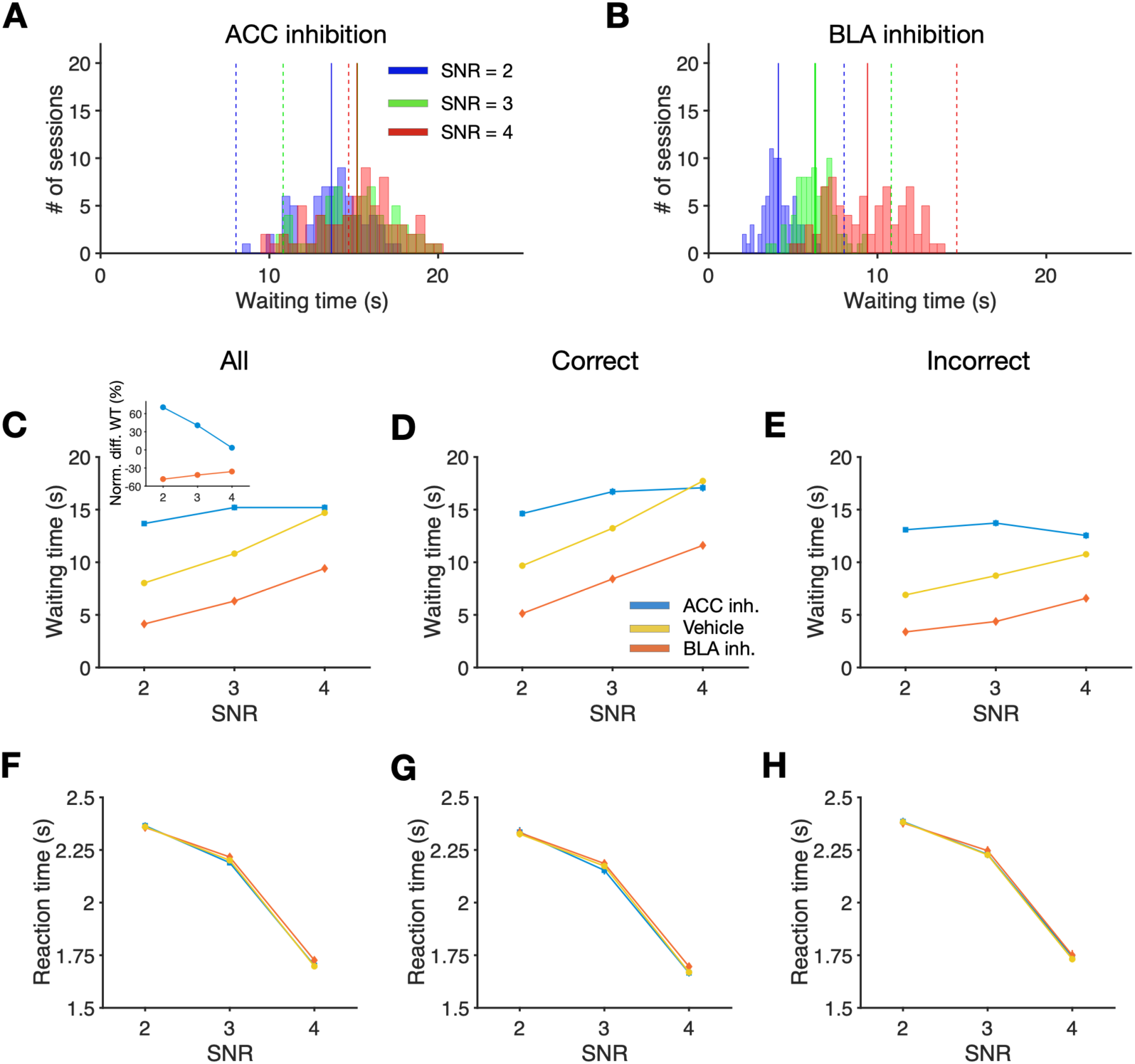
ACC and BLA inhibition result in opposing changes in waiting time, but no change in reaction times. **(A)** Waiting time increases following ACC inhibition. Plotted are the distributions of the waiting time separately for each SNR, following ACC inhibition. Solid lines show the median of each distribution. Dashed lines show the median of the same condition but following vehicle administration as is shown in Fig. 2A. **(B)** Waiting time decreases following BLA inhibition. Same as in panel A but for sessions following inhibition of BLA. **(C)** Inhibition of ACC renders waiting times insensitive to the strength of sensory signal, whereas BLA inhibition shift waiting times. Plotted is the waiting time for all trials as a function of SNR following vehicle administration (yellow), inhibition of BLA (orange), and inhibition of ACC (blue). (**D-E**) Same as in panel C but only on trials in which a correct (D) or incorrect (E) response was made. Error bars show the S.E.M. over sessions (typically smaller than the symbols). (**F**) Reaction time is unaffected by inhibition of either ACC or BLA. Plotted is the reaction time for all trials as a function of SNR following vehicle administration (yellow), inhibition of BLA (orange), and inhibition of ACC (blue). (**G-H**) Same as in panel F but only on trials in which a correct (G) or incorrect (H) response was made.

Averaging over different contrast levels, we found that the probability of making a correct response was larger when SNR was larger (GLM; *F*(3,437) = 106, *p* = 1.1 × 10^−51^, adjusted *R*^2^ = 0.418; ratio: *β* = 0.054, *p* = 1.3 × 10^−29^; *p*(correct) = 0.69 ± 0.06, 0.74 ± 0.05, and 0.80 ± 0.04 for SNR of 2, 3, and 4, respectively). In addition, the number of re-initiations decreased as SNR increased (GLM; *F*(3,437) = 17.7, *p* = 7.6 × 10^−11^, adjusted *R*^2^ = 0.102; SNR: *β* = −0.01, *p* = 4.36 × 10^−6^; fraction of re-initiated trials = 0.17 ± 0.02, 0.16 ± 0.03, and 0.15 ± 0.02 for SNR of 2, 3, and 4, respectively). Finally, the distribution of waiting time generally followed that of reward delivery (**Supplementary Figure 4**). These results illustrate that rats learned the task and used visual stimuli to make a choice and for post-choice wagering.

To show that waiting time to re-initiate a trial met several of the criteria for confidence readout, we analyzed this measure along with reaction time to make a response (i.e., the time between stimulus onset and nosepoke) during the control sessions. We found that waiting times were longer on trials with a larger SNR (GLM; *F*(7,874) = 221, *p* = 2.5 × 10^−188^, adjusted *R*^2^ = 0.636; SNR: *β* = 1.91, *p* = 1.2 × 10^−16^; **Figure 2A**). This effect has previously been reported for confidence reports in human and nonhuman primates (Koizumi et al., 2015; Miyamoto et al., 2017; Rounis et al., 2010), but not in rodents. In contrast, reaction time decreased with increased SNR (GLM; *F*(7,874) = 221, *p* = 3.04 × 10^−216^, adjusted *R*^2^ = 0.686; SNR: *β* = −0.32, *p* = 1.27 × 10^−78^; **Figure 2B**). Additionally, waiting times were negatively correlated with reaction times on a trial-by-trial basis (GLM; *F*(1,880) = 318, *p* = 4.98 × 10^−61^, adjusted *R*^2^ = 0.265; reaction time: *β* = −7, *p* = 4.97 × 10^−61^; **Figure 2C**). This negative correlation between post-decision wagering and reaction time involving visual stimuli has also been reported in primates, but not in rodents.

To examine the relationship between accuracy and time wagering, we computed waiting time separately for correct and incorrect responses. We found that rats waited significantly longer following correct relative to incorrect responses, or trial type (diff(mean)=4.74; GLM: *F*(3,878) = 123, *p* = 1.31 × 10^−66^, adjusted *R*^2^ = 0.294; trial type: *β* = 4.57, *p* = 2.46 × 10^−33^; **Figure 2D**). Consistent with this result, on trials with a re-initiation, rats discriminated more accurately when waiting times were longer for a given SNR (Pearson correlation; SNR=2: *r* = 0.017, *p* = 0.83; SNR = 3, *r* = 0.19, *p* = 0.017; SNR = 4, *r* = 0.19, *p* = 0.02; **Figure 2A** inset). In addition, waiting times on both correct and incorrect responses increased with larger SNR (GLM: Correct; *F*(3,437) = 189, *p* = 1.95 × 10^−78^, adjusted *R*^2^ = 0.561; SNR: *β* = 3.74, *p* = 1.40 × 10^−41^; Incorrect; *F*(3,437) = 68.8, *p* = 1.77 × 10^−36^, adjusted *R*^2^ = 0.316; SNR: *β* = 1.91, *p* = 2.33 × 10^−19^). Finally, the normalized difference in waiting times between correct and incorrect responses changed strongly (30-50%) for different SNR values (**Figure 2D** inset). These results demonstrate that not only is waiting time sensitive to the strength of the visual information, but it also reflects rats’ accuracy in discrimination.

Compatible with previous findings, we also found that reaction times decreased with larger SNR and were faster for correct responses relative to incorrect responses/discrimination (GLM; *F*(3,878) = 2.56, *p* = 0.05, adjusted *R*^2^ = 0.005; SNR: *β* = −0.069, *p* = 0.03; **Figure 2E**). The normalized difference in reaction times between correct and incorrect responses, however, changed only between 1-5% for different SNR values compared to 30-50% for waiting times (**Figure 2E** inset). In addition, unlike waiting time, there was no significant correlation between performance and reaction time on a session-by-session basis (Pearson correlation; SNR=2, *r* = −0.1, *p* = 0.2; SNR=3, *r* = 0.13, *p* = 0.1; SNR =4, *r* = 0.01, *p* = 0.82; **Figure 2B**). Importantly, we found similar results when we performed all above analyses for each value of contrast separately (**Supplementary Figure 2**). Moreover, independently of SNR and contrast levels, waiting time increased with discrimination performance (**Supplementary Figure 1B**).

Together, our results illustrate that waiting time reflects confidence in a perceptual discrimination with much higher fidelity than that of reaction time, to include the proportional nature of confidence and accuracy. Our findings thus extend previous observations in primates (Kiani & Shadlen, 2009) to rodents, and suggest that waiting time in our paradigm can also serve as a proxy for decision confidence (Lak et al., 2014).

### Dissociable contributions of BLA and ACC to time wagering

We expressed Gi-coupled DREADD receptors in projection neurons of ACC and BLA (**Figure 1C**). After allowing time for transduction, we injected rats with CNO prior to a subset of testing sessions to inhibit these brain regions, using a within-subject design. In addition, to confirm the effect of CNO using *ex vivo* electrophysiological recording, we prepared a separate group of rats (*n* = 3) with ACC DREADDs using identical procedures. We found a significant reduction in field potential after CNO application only in the transfected slices (**Supplementary Figure 5**).

For rats performing the main experiments, we observed significant interaction of drug condition (vehicle and CNO), targeted brain region (BLA or ACC), and SNR on waiting time when combining correct and incorrect responses (GLM; *F*(15,1742) = 327, *p* = 10^−17^, adjusted *R*^2^ = 0.735; drug × region × ratio: *β* = 1.84, *p* = 7.4 × 10^−6^). We found no significant effect of the targeted brain region (BLA vs. ACC) in sessions following vehicle administration on waiting time (GLM; *F*(7,874) = 221, *p* = 2.5 × 10^−188^, adjusted *R*^2^ = 0.636; region: *β* = −0.93, *p* = 0.335). In contrast, following CNO administration, we observed a significant effect of the targeted brain region on waiting time (GLM; *F*(7,868) = 518, *p* = 6.36 × 10^−305^, adjusted *R*^2^ = 0.805; region: *β* = 1.66, *p* = 5.11 × 10^−55^; **Figure 3A,B**).

An analysis of waiting time for different SNR values averaged across all trial types (correct and incorrect) revealed a significant drug × SNR × brain region interaction (GLM; *F*(7,1750) = 282, *p* = 7.07 × 10^−282^, adjusted *R*^2^ = 0.529; three-way interaction: *β* = 1.66, *p* = 1.94 × 10^−5^). When the trial type (correct vs incorrect) was included as a within-subject factor, there was similarly a significant trial type × drug × brain region interaction (GLM; *F*(15,1742) = 327, *p* = 10^−17^, adjusted *R*^2^ = 0.735, three-way interaction: *β* = 1.84, *p* = 7.40 × 10^−6^). These results show that inhibition of ACC or BLA affect rats’ willingness to wait depending on SNR (perceptual uncertainty) as well as based on whether their response was correct or incorrect.

Given these interaction effects, we then measured the influence of inhibition of ACC and BLA on waiting time separately for each value of SNR. We found that overall, inhibition of ACC significantly increased waiting time compared to vehicle (diff(mean)=3.45 sec; GLM; *F*(1,1288) = 204, *p* = 4.98 × 10^−43^, adjusted *R*^2^ = 0.136; drug: *β* = 3.45, *p* = 4.98 × 10^−43^; **Figure 3A,C**). In contrast, inhibition of BLA significantly reduced rats’ overall willingness to wait before re-initiation of a new trial (diff(mean)=-4.59sec; GLM; *F*(1,1348) = 388, *p* = 4.87 × 10^−76^, adjusted *R*^2^ = 0.223; drug: *β* = −4.58, *p* = 4.86 × 10^−76^; **Figure 3B,C)**. Importantly, we found a significant interaction of SNR by drug condition (CNO vs. vehicle administration) for ACC (GLM; SNR × drug: *β* = −2.19, *p* = 1.78 × 10^−10^), indicating that the observed increase in waiting time due to ACC inhibition depended on SNR. In contrast, the decrease in waiting time due to BLA inhibition was not SNR-specific (GLM; SNR × drug: *β* = −0.3434, *p* = 0.16), indicating that BLA inhibition increased sensitivity to delays (or equivalently increased impulsivity), as has been reported before (Winstanley, Theobald, Cardinal, & Robbins, 2004). Finally, we found similar results when we performed our analyses for each value of contrast separately (**Supplementary Figure 3A-C**).

Despite strong, dissociable effects on waiting time, inhibition of ACC and BLA did not change the overall task performance, discrimination accuracy, response bias, or reaction time. First, probability of correct response was not significantly different between vehicle and CNO administration (GLM; *F*(3,289) = 2.77, *p* = 0.04, adjusted *R*^2^ = 0.017; drug: *β* = 0.003, *p* = 0.57; **Figure 4A**). Second, we computed discrimination performance or *d*′ and found this measure was also not significantly different between vehicle and CNO administration (GLM; *F*(7,82) = 40.7, *p* = 4.19 × 10^−24^, adjusted *R*^2^ = 0.757; drug: *β* = −0.26, *p* = 0.21; **Figure 4B**). Importantly, we observed a strong and significant correlation between discrimination accuracy *d*′ following CNO administration and following vehicle administration in rats with DREADDs expressed either in ACC (Pearson correlation; *r* = 0.847, *p* = 1.25 × 10^−6^) or BLA (Pearson correlation; *r* = 0.767, *p* = 1.21 × 10^−5^). Moreover, contrary to the significant effect of SNR on discrimination accuracy *d*′ (GLM; *F*(7,82) = 40.7, *p* = 4.19 × 10^−24^, adjusted *R*^2^ = 0.757; ratio: *β* = 0.38, *p* = 2.38 × 10^−11^), we found no significant main effect or interaction of drug condition (CNO vs. vehicle administration) and targeted brain region (ACC vs. BLA) on *d*′ (GLM; drug: *β* = −0.26, *p* = 0.21; region: *β* = 0.04, *p* = 0.84), indicating that perceptual discrimination was not affected by ACC or BLA inhibition. Third, we found no significant effect of drug condition (CNO vs. vehicle) and targeted brain region on the decision criterion (i.e., the response bias; (Macmillan & Creelman, 1997); GLM; *F*(3,86) = 1.62, *p* = 0.19, adjusted *R*^2^ = 0.02; drug: *β* = −0.03, *p* = 0.23; region: *β* = −0.007, *p* = 0.74). Finally, ACC and BLA inhibition failed to affect task engagement and perceptual processing speed as evidenced by the lack of change in the distributions of reaction time (GLM; *F*(15,1742) = 234, *p* = 10^−16^, adjusted *R*^2^ = 0.665; drug: *β* = −0.01, *p* = 0.81; region: *β* = 0.001, *p* = 0.98; **Figure 3F**), and these responses also did not differ by trial type (correct vs. incorrect; Correct: GLM; *F*(7,871) = 260, *p* = 2.01 × 10^−208^, adjusted *R*^2^ = 0.674; ratio: *β* = −0.33, *p* = 1.01 × 10^−81^; Incorrect: GLM; *F*(7,871) = 236, *p* = 3.44 × 10^−196^, adjusted *R*^2^ = 0.652; ratio: *β* = −0.32, *p* = 3.14 × 10^−72^; **Figure 3G,H**). This pattern also held for each value of contrast separately (**Supplementary Figure 3D-F**). Together these findings demonstrate that the observed effects of ACC and BLA inhibition on waiting times were not attributable to changes in decision-making processes related to visual discrimination.

**Figure 4.**
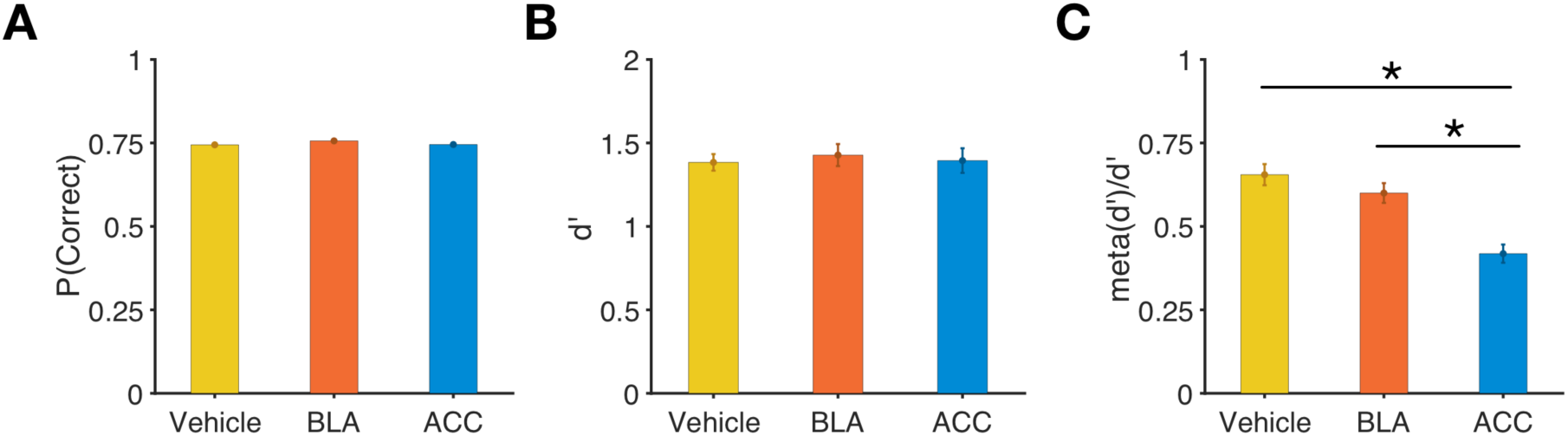
Discrimination performance is intact following inhibition of either ACC or BLA, but metacognitive efficiency is decreased following inhibition of ACC only. **(A)** Plotted is the probability of correct responses for the sessions following vehicle administration (yellow), sessions following BLA inhibition (orange), and sessions following inhibition of ACC (blue). Error bars show S.E.M. **(B)** Plotted is discrimination performance, *d*′ in the three conditions defined in panel A. **(C)** Plotted is the metacognitive efficiency (meta-*d*′/*d*′) in the three conditions defined in panel A. (*) *p* < 0.05.

Finally, in additional control conditions, we also tested whether the presence of active virus was essential for the observed changes, and whether vehicle administration alone could cause changes in behavior. To do so, we measured behavioral responses in the rats with expressed DREADDs but without the administration of vehicle (no-injection control prior to reversal) and in rats with null virus and compared them with those under vehicle administration (i.e., the main control condition). We found that waiting time and reaction time did not differ between vehicle administration and no-injection control (see Supplementary analyses and **Supplementary Figure 6**). In addition, we found that the observed effects of CNO depended on the presence of active virus (Supplementary analyses).

Together, these results suggest that ACC contributes to the computations and transmission of confidence to influence post-decision behavior. In contrast, BLA mainly increases waiting time independently of perceptual uncertainty perhaps by controlling impulsive behavior during choice under uncertainty.

### ACC-specific role in evaluation for confidence report

Although BLA inhibition decreased waiting time, this measure still increased with greater SNR (GLM; *F*(3,464) = 324, *p* = 2.32 × 10^−113^, adjusted *R*^2^ = 0.675; SNR: *β* = 1.59, *p* = 5.44 × 10^−23^; **Figure 3C**) similar to the behavioral pattern observed under vehicle administration (GLM; *F*(3,878) = 500, *p* = 1.63 × 10^−189^, adjusted *R*^2^ = 0.63; SNR: *β* = 1.92, *p* = 3.16 × 10^−32^). This suggests that BLA contributes to shifting the influence of confidence on post-decision processes making the animal more patient irrespective of their confidence. In contrast, ACC inhibition rendered waiting time mainly insensitive to SNR (GLM; *F*(3,404) = 55.1, *p* = 7.17 × 10^−30^, adjusted *R*^2^ = 0.28; SNR: *β* = −0.27, *p* = 0.22; **Figure 3C**), with a significant effect of SNR on correct trials (GLM; *F*(1,202) = 26.5, *p* = 6.09 × 10^−7^, adjusted *R*^2^ = 0.11; ratio: *β* = 1.22, *p* = 6.09 × 10^−7^; **Figure 3D**), but not for incorrect trials (GLM; *F*(1,202) = 1.65, *p* = 0.2, adjusted *R*^2^ = 0.0032; ratio: *β* = −0.27, *p* = 0.2; **Figure 3E**). Interestingly, ACC inhibition renders waiting time even insensitive to the contrast level of visual stimuli such that for higher contrast and higher SNR, waiting time following inhibition dropped below the control condition (**Supplementary Figure 3A-C**). These findings suggest that ACC is involved in modifying visual uncertainty, perhaps via gain modulation, in order to compute perceptual uncertainty and to influence post-decision processes based on the latter.

To further test this, we computed metacognitive efficiency (meta-*d*′/*d*′; see Methods for details), that assesses how well waiting time tracks discrimination performance (*d*′) across trials (Maniscalco & Lau, 2012), or equivalently, the trial-by-trial correspondence of accuracy and waiting time. We found that this measure was significantly reduced following ACC inhibition compared to vehicle administration (Wilcoxon rank-sum test; *p* = 1.037 × 10^−7^; **Figure 4C**) but remained intact following BLA inhibition (Wilcoxon rank-sum test; *p* = 0.11). Nonetheless, metacognitive efficiency was larger than zero in both vehicle and CNO administration conditions (t-test; vehicle: *t*(134) = 20.65, *p* = 1.763 × 10^−43^; ACC inhibition: *t*(62) = 15.33, *p* = 9.128 × 10^−23^; BLA inhibition: *t*(71) = 20.24, *p* = 3.304 × 10^−31^) and significantly less than 1 (t-test; vehicle: *t*(134) = −10.88, *p* = 3.751 × 10^−20^; ACC inhibition: *t*(62) = −21.3, *p* = 3.172 × 10^−30^; BLA inhibition: *t*(71) = −13.49, *p* = 2.787 × 10^−21^). Consistent with these results, we found that the trial-by-trial correlation between waiting time and reaction time was weaker following ACC inhibition compared to BLA inhibition (**Supplementary Figure 7**).

Collectively, these results suggest that whereas inhibition of BLA decreases waiting time, this effect is most likely due to the general delay aversion or an increase in impulsive choice, because rats are still able to appropriately scale their waiting times according to performance and trial difficulty. In contrast, inhibition of the ACC renders rats’ waiting times relatively insensitive to discrimination accuracy (*d*′) and SNR, suggesting that this region meaningfully participates in estimating the reliability of visual stimuli and consequently, computing and reporting confidence. Taken together with the results we provide in the previous section, we show that here in rats we are able to interfere with and dissociate first order (discrimination performance) from second order (metacognition) processes, as has been done in nonhuman and human primates (Koizumi et al., 2015; Maniscalco, Peters, & Lau, 2016; Miyamoto, Setsuie, Osada, & Miyashita, 2018; Odegaard et al., 2018).

### Confidence enhances reversal learning

Following learning of the task and the stimulus-response rule, rats were randomly assigned into a high confidence (HC) or low confidence (LC) condition, and subsequently experienced a reversal in the stimulus-response rule. Unlike in the initial task, upon reversal, rats were not permitted to re-initiate the trial while waiting for a possible reward delivery. Correct responses, now under a reversed stimulus-response rule, were reinforced probabilistically as before (70% of the time). We selected the two parameters of the stimulus (SNR and contrast level) for each rat from a pair of matched-performance (i.e., matched discrimination accuracy, or *d*′) but different average waiting time. This was only possible because as we illustrated earlier, SNR and contrast allowed us to modulate performance and waiting time independently. As we show below, although we used different visual stimuli across HC and LC conditions, there was no systematic difference in performance accuracy and instead, there was difference only in confidence levels measured by the waiting time before the reversal. Importantly, we calculated *d*′ and confidence using only the data from sessions that were not preceded by injections (i.e., no-injection control prior to reversal) in order to assign rats to HC and LC conditions.

We performed several analyses to ensure that the only difference between HC and LC conditions was the confidence reported via waiting time. First, we found that *d*′ for HC and LC conditions were not significantly different for each of the stimuli that was administered after reversal, i.e., the contrast-SNR pairs that were chosen for reversal were not associated with different *d*′ before reversal (Stepwise GLM; *F*(3,11) = 2.46, *p* = 0.11, adjusted *R*^2^ = 0.238; confidence: *β* = 0.43, *p* = 0.32). Second, *d*′ for HC and LC conditions were not significantly different across all contrast-SNR pairs (GLM; *F*(15,119) = 10.6, *p* = 8.2 × 10^−16^, adjusted *R*^2^ = 0.52; confidence: *β* = −0.14, *p* = 0.92). Third, metacognitive efficiency (meta-*d*′/*d*′) across HC and LC conditions was not significantly different for the specific contrast-SNR pair that was used after reversal (Stepwise GLM; *F*(1,13) = 0.62, *p* = 0.44, adjusted *R*^2^ = −0.03; confidence: *β* = −0.1, *p* = 0.44). Fourth, rats in both HC and LC conditions acquired equal amount of reward in the no-injection control session for the specific pair of contrast and SNR values (Stepwise GLM; *F*(3,148) = 3.57, *p* = 0.01, adjusted *R*^2^ = 0.05; confidence: *β* = −0.2, *p* = 0.08). Finally, we found that HC and LC conditions were different in waiting time, reflecting confidence, for the specific contrast-SNR pair that was used after reversal (Stepwise GLM; *F*(5,146) = 53.4, *p* = 2.7 × 10^−31^, adjusted *R*^2^ = 0.634; confidence: *β* = −28.3, *p* = 2.81 × 10^−9^). Together, these results illustrate that the only difference between HC and LC conditions was the confidence.

To assess the effect of perceptual uncertainty on learning, we analyzed the probability of correct response for both HC and LC conditions for rats receiving vehicle or CNO injections, and for ACC or BLA as the targeted brain region. We observed significant interactions of drug by confidence level (GLM; *F*(7,169) = 226, *p* = 2.95 × 10^82^, adjusted *R*^2^ = 0.899; drug × confidence: *β* = −0.22, *p* = 3.71 × 10^−7^ as well as drug by trial bin (*β* = −0.009, *p* = 0.0001). There was also a significant main effect of trial bin (*β* = 0.036, *p* = 1.13 × 10^−44^), illustrating that all rats were able to learn the new stimulus-response rule. Similarly, there was a significant main effect of confidence on learning (*β* = 0.15, *p* = 3.4 × 10^−6^).

To identify how learning and choice strategies were affected by confidence, we first compared learning between HC and LC conditions following vehicle administration. We found that rats in the HC condition performed better than the rats in the LC condition following vehicle administration (diff(mean)=0.1063; GLM; *F*(3,75) = 262, *p* = 1.13 × 10^−39^, adjusted *R*^2^ = 0.91; confidence: *β* = 0.151, *p* = 2.71 × 10^−6^). This improvement in performance was due to faster learning in the HC compared to LC condition (Chi-square test of ratio; *p* = 5.87 × 10^−26^; **Figure 5A inset**) and the steady state of performance was not affected by confidence (Chi-square test of ratio; *p* = 0.27).

**Figure 5.**
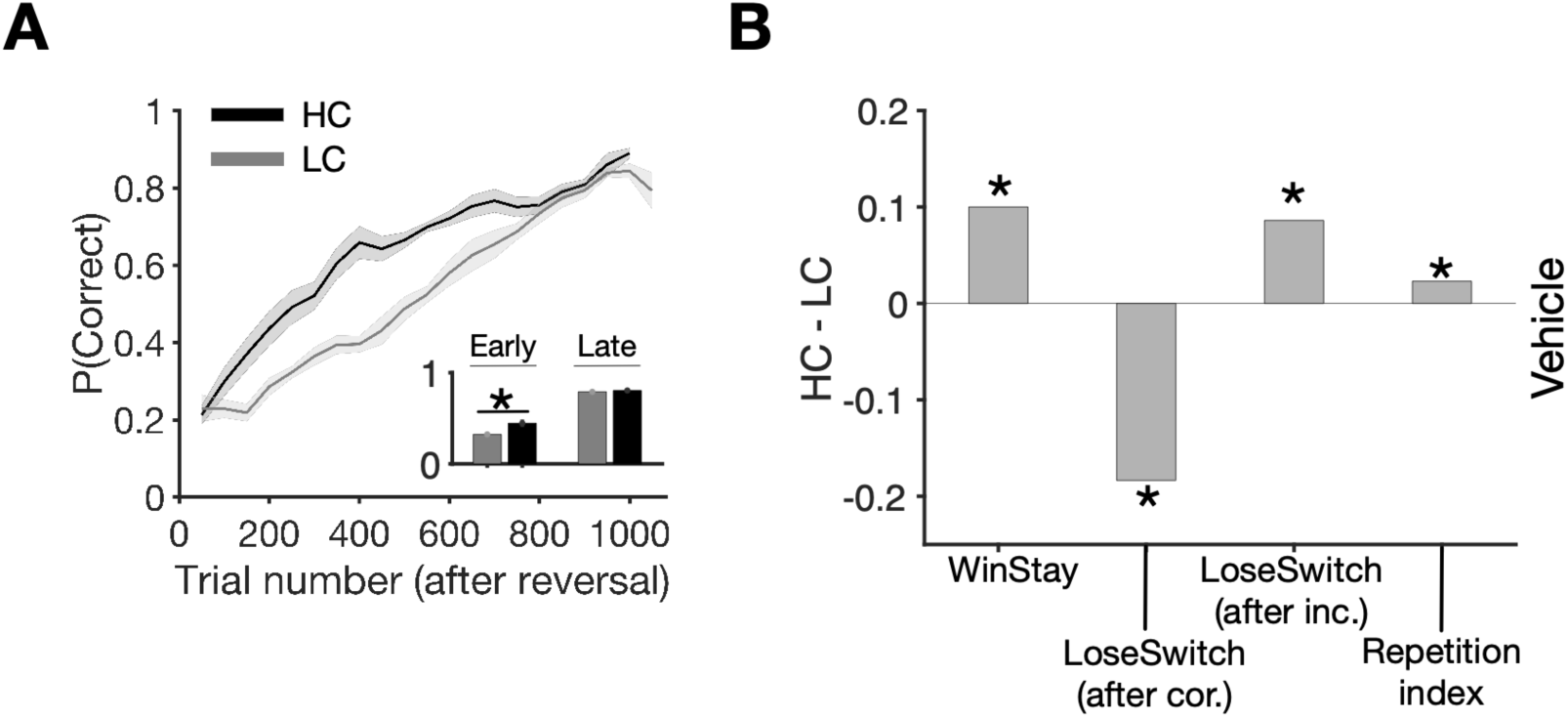
Influence of decision confidence on subsequent reversal learning. **(A)** Learning curves showing performance (probability of correct response) following reversal for rats prepared with DREADDs after vehicle administration. Plots shows the performance across all rats in high-confidence (HC; black) and low-confidence (LC; gray) conditions averaged over a sliding window of 100 trials. The inset shows the average performance in early (first half of the trials) and late (last quarter of the trials) trials, demonstrating that higher confidence improved the rate of learning but did not change the steady state. (*) indicates a significant difference in median between the two conditions (Chi-square test of ratio, *p* < 0.05). **(B)** The influence of confidence on learning strategies and perseveration. Plotted are the difference in the proportions of Win-Stay, Lose-Switch following correct but unrewarded responses, Lose-Switch after incorrect responses, and rule-based repetition index between HC and LC conditions in the first half of trials after the reversal.

The observed faster learning occurred simultaneously with an increase in selection of the correct stimulus-response rule following selection of this rule and being rewarded on the preceding trial (Win-Stay; Permutation test; *p* = 9.7 × 10^−10^; **Figure 5B**). In addition, animals increased their tendency to switch from the incorrect to correct stimulus-response rule following unrewarded trials when the response on the preceding trial was incorrect (Lose-Switch after incorrect; Permutation test; *p* = 0.046). The improvement in learning due to higher confidence was also accompanied by a decrease in switch from the correct to incorrect stimulus-response rule when the response on the preceding trial was correct but not rewarded (Lose-Switch after correct; note that 30% of correct responses were not rewarded by design; Permutation test; *p* = 3.3 × 10^−4^). We also compared the tendency of the animals to repeat the same stimulus-response rule as in the previous trial beyond what is expected by chance, measured by the rule-based repetition index (RRI; (Soltani, Noudoost, & Moore, 2013); see Methods). We found that RRI was larger for the HC relative to LC condition (Permutation test; *p* = 0.029), indicating that animals were more consistent/persistent in their behavior (following a specific rule) under higher confidence. Together, these results suggest that confidence can improve learning strategies from *all possible outcomes* and moreover, can increase consistency in following learned stimulus-response rules. To our knowledge, the observed enhancing effect of perceptual confidence on learning has been reported in humans (Guggenmos et al., 2016) but not in rodents, and the effects on rule consistency are novel.

### Both BLA and ACC support reversal learning

We next compared overall learning across different DREADDS inhibition conditions. In contrast to the control conditions, the overall performance over time in the LC condition was significantly better than in the HC condition following CNO treatment (diff(mean)=0.0456; GLM; *F*(3,94) = 269, *p* = 5.62 × 10^−46^, adjusted *R*^2^ = 0.892; confidence: *β* = −0.0719, *p* = 0.01). Furthermore, the reduction in performance was not significantly different between ACC and BLA as the targeted region (GLM; *F*(7,90) = 114, *p* = 5.78 × 10^−42^, adjusted *R*^2^ = 0.891, region: *β* = −0.035, *p* = 0.396; **Figure 6A,C**). To estimate the rate of learning, we fit rats’ learning curves after the reversal using an exponential function (see Methods for details). We observed that the exponent or the learning parameter (ɑ; which reflects the rate of learning) was significantly different between CNO and vehicle administration sessions in the HC condition (confidence interval; ACC; CNO, [15.2,15.78], vehicle, [9.42,10.04]; BLA; CNO, [13.28,16.20], vehicle, [8.23,9.23]). However, there was no significant difference in the learning parameter between sessions following CNO and vehicle administration in the LC condition (confidence interval; ACC; CNO, [12.52,13.75], vehicle, [12.36,15.18]; BLA; CNO, [12.78,14.12], vehicle, [12.63,14.33]). Therefore, following inhibition of either ACC or BLA, the rate of learning decreased in the HC but not the LC condition. Together, these results indicate that whereas rats could still learn a new stimulus-response rule after the inhibition of the ACC or BLA, these brain regions contribute to an *enhancing* effect (i.e. the use) of perceptual confidence on learning despite matched *d*′ across HC and LC conditions.

**Figure 6.**
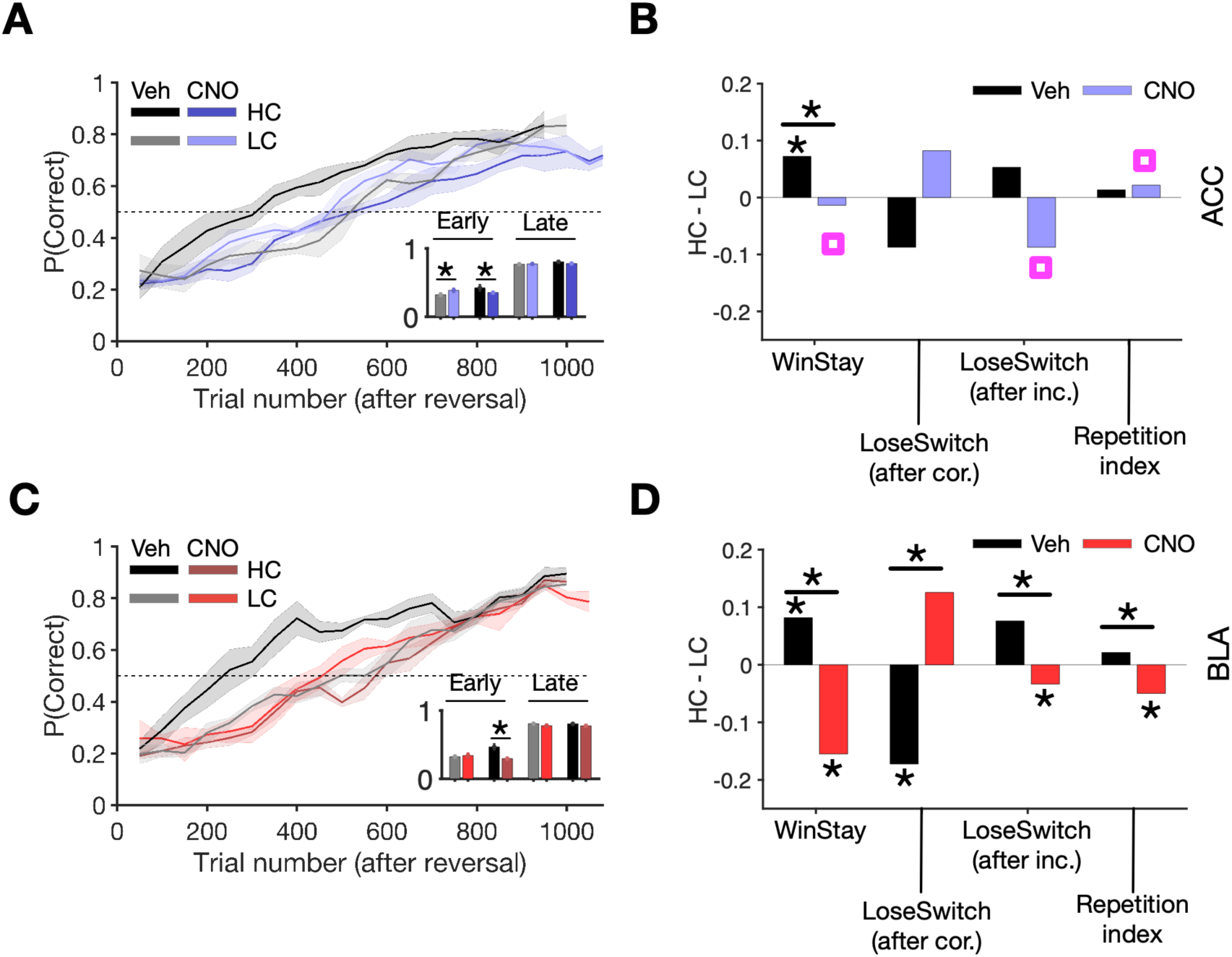
ACC and BLA inhibition differentially modulate the effects of perceptual uncertainty on learning strategies and perseveration. **(A)** Learning curves (probability of correct response) after a reversal for rats prepared with ACC DREADDs following CNO (blue) or vehicle (Veh; black) administration. Plot shows the performance across all rats averaged over a sliding window of 100 trials for high-confidence (HC) and low-confidence (LC) conditions. The inset shows the average performance in early (first half of the trials) and late (last quarter of the trials) trials, demonstrating that either perceptual uncertainty or ACC inhibition decrease the rate of learning. Following ACC inhibition, rats eventually reach a similar performance level compared to the control condition (vehicle administration). (*) indicates a significant difference in median between the two conditions (Chi-square test of ratio, *p* < 0.05). (**B**) Win-Stay, Lose-Switch following correct response, Lose-Switch after incorrect response, and rule-based repetition index for the HC and LC conditions in ACC DREADDs during the first half of trials after the reversal. ACC inhibition only removes the benefit of confidence on Win-Stay but weakens the effect of confidence on learning from negative feedback or consistency in rule selection. (*) indicates a median significantly different from zero or a significant difference in median between the two conditions (Permutation test, Bonferroni corrected, *p* < 0.01). Magenta squares indicate a significance difference between ACC and BLA inhibition (Permutation test, *p* < 0.05). **(C)** Learning curves (probability of correct response) after a reversal for rats prepared with BLA DREADDs following CNO (red) or vehicle (black) administration. BLA inhibition decreases the rate of learning but eventually rats reach a similar performance level compared to the vehicle administration condition. (**D**) The same as in panel B but for BLA DREADDs. Unlike ACC inhibition, BLA inhibition reverses the benefits of confidence on all learning strategies and consistency in rule selection.

### BLA and ACC use confidence information for learning strategy

Results reported above suggest that both types of DREADDs inhibition (in ACC or BLA) removed the benefit of confidence on learning. To better reveal similarities and differences between the effects of ACC and BLA inhibition on learning and their dependence on confidence, we computed changes in the learning strategies and consistency in following stimulus-response rules after ACC and BLA inhibition, separately for rats in the LC and HC conditions.

First, we examined the effect of positive feedback on the tendency to stay on the correct stimulus-response rule (Win-Stay). We found that inhibition of either ACC or BLA resulted in a reduction in the difference in Win-Stay between HC and LC conditions (Permutation test; ACC: *p* = 5.43 × 10^−4^; BLA: *p* = 5.83 × 10^−12^; **Figure 6B,D**) but this reduction was stronger for BLA inhibition (Permutation test; *p* = 0.0048). This suggests that both ACC and BLA inhibition attenuate reversal learning by reducing the tendency to repeat a rewarded stimulus-response rule due to higher confidence. Therefore, although both ACC and BLA contribute to mediating the effect of confidence on learning from positive feedback, the stronger attenuation and reversal of this effect following BLA inhibition illustrates a more prominent role for BLA in learning under uncertainty.

Secondly, we analyzed the effect of negative feedback on switching from the previous stimulus-response rule, separately for when the previous rule was correct (Lose-Switch after correct) and when the previous rule was incorrect (Lose-Switch after incorrect). We examined these two types of trials separately because as we showed above, confidence has differential effects on these trials (**Figure 5B**). We observed that both ACC and BLA inhibition reversed the effect of confidence (i.e., difference between HC and LC conditions) on learning from negative feedback; however, this was only significant for BLA (Permutation test; ACC Lose-Switch after correct: *p* = 0.23; ACC Lose-Switch after incorrect: *p* = 0.44; BLA Lose-Switch after correct: p=7.4*10^−4^; BLA Lose-Switch after incorrect: p=0.0044; **Figure 6B,D)**. In addition, the effect of confidence on Lose-Switch after an incorrect response became more negative after ACC inhibition compared to BLA inhibition (Permutation test; *p* = 0.0051), indicating a stronger role for ACC in learning from negative feedback when the response was incorrect. Together, these results suggest that BLA has a more pronounced role in mediating the effect of confidence in learning from positive feedback whereas ACC is more involved in mediating the effect of confidence in learning from negative feedback.

Finally, we evaluated the consistency in using learned stimulus-response rules using the RRI. We found that BLA but not ACC inhibition reversed the effect of confidence on RRI (Permutation test; ACC: *p* = 0.36; BLA: *p* = 4.6 × 10^−4^; **Figure 6B,D**), and this effect was more negative after BLA than ACC inhibition (Permutation test; *p* = 4.4 × 10^−4^). This indicates that BLA, but not ACC, is important for mediating the effect of confidence in consistently using learned stimulus-response rules under perceptual uncertainty. Importantly, there was no significant difference between the effects of confidence on all aforementioned strategies in sessions following vehicle administration based on the targeted region (Permutation test; Win-Stay: *p* = 0.31; Lose-Switch after correct: *p* = 0.10; Lose-Switch after incorrect: *p* = 0.35; RI: *p* = 0.36; **Figure 6B,D**).

Together, these results reveal dissociable effects of ACC and BLA inhibition on learning under uncertainty. Importantly, only BLA inhibition consistently reversed the benefit of confidence on learning from both positive and negative feedback. This suggests that BLA is directly involved in confidence-dependent learning (and not estimation) because BLA inhibition only shifts confidence readout with respect to perceptual uncertainty, as shown earlier. Different from BLA effects, ACC has a more specific role in supporting learning, mainly from negative feedback following an incorrect response and to lesser extent from positive feedback (Win-Stay), perhaps by making confidence computation sensitive to the level of perceptual uncertainty, as suggested by our results on confidence readout. In addition, the consistency in using learned stimulus-response rules did not differ by confidence condition following ACC inhibition, whereas BLA inhibition made rats less likely to apply the learned rules under higher confidence.

## Discussion

We examined the causal roles of ACC and BLA in confidence report and learning under perceptual uncertainty. We studied learning under uncertainty by training rats to report the orientation of ambiguous visual stimuli based on a learned stimulus-response rule and read out their confidence in choice using a time wagering task. Decision opt-out and wagering tasks have been previously used to assess confidence in rats (Foote & Crystal, 2007; Lak et al., 2014), showing that rats exhibit similar behavior to those of humans with monetary rewards (Persaud, McLeod, & Cowey, 2007). We observed that rats are willing to tolerate larger delays to outcomes after faster and easier perceptual decisions involving more salient stimuli. We showed that ACC is required for appropriate waiting according to the uncertainty of the visual stimulus and ensuing choice. Following ACC inhibition, post-decision waiting times were less sensitive to the strength of the visual evidence, and accuracy tracked less well with these waiting times on a trial-by-trial basis. In contrast, inhibition of BLA decreased rats’ willingness to wait overall, regardless of the strength of the visual information and decision difficulty.

It has been proposed that confidence in a decision not only affects our choices but also influences how we learn (Guggenmos et al., 2016; Kiani & Shadlen, 2009). However, the effect of high confidence on reinforcement learning had not been explored directly in any animal model. We found that high confidence in a perceptual decision can boost subsequent reversal learning of stimulus-response rules using reward feedback, even when we controlled for signal processing capacity, (i.e., task performance). Critically, all rats were able to learn new reward contingencies upon the change in stimulus-response mapping, but the learning was faster in the group of rats that had higher confidence at the onset of reversal. We show that the BLA and ACC are both required for the enhancement of learning by perceptual certainty or confidence.

### ACC in perceptual decision making

In rats, perceptual metacognition has been previously assessed within olfactory and auditory (Foote & Crystal, 2007; Kepecs et al., 2008; Lak et al., 2014) but not visual modalities. These studies have revealed a role of orbitofrontal cortex (OFC); for example, it has been shown that activity in the rat OFC reflects the degree of uncertainty in decisions based on olfactory information during reward anticipation (Kepecs et al., 2008). Similar to our results for the ACC, inhibition of OFC impairs behavioral adjustments to decision confidence, but not perceptual choices based on conflicting evidence themselves (Lak et al., 2014). However, there are several important differences, not just similarities, between previous and present studies. First, Lak et al. (2014) showed that waiting time increased for correct trials and decreased for incorrect trials, whereas we show that waiting time increased for both correct and incorrect trials, across all contrast levels. This relationship, however, depended on the ACC. Specifically, following ACC inhibition, waiting time became insensitive to SNR especially on trials with an incorrect response. Nevertheless, here, we show that the ACC plays a similar role to the OFC but in visual information processing. That is, the ACC may guide commitment to and persistence with the current behavior based on the quality of visual evidence that led to the decision. These similarities offer interesting possibilities for the frontocortical mechanisms of confidence estimation and suggest there may not be a subregional specialization for this process (Hunt & Hayden, 2017; Yoo & Hayden, 2018). Consequently, future research should be directed at uncovering the constraints (and if there is differential involvement that may be revealed over different timescales) for the ACC and OFC in decisional confidence.

We show here that unlike BLA inhibition, ACC inhibition renders confidence readout rather insensitive to both attributes of visual stimuli (SNR and contrast), suggesting that ACC “gain” modulates visual uncertainty computed in visual areas to determine perceptual uncertainty and post-decision processes. Anatomically, the ACC is densely interconnected with visual cortices in rodents (Vogt & Miller, 1983; Vogt & Paxinos, 2014), particularly the more rostral aspect of ACC in rat as we have targeted here (Vogt & Paxinos, 2014). Furthermore, this brain region is well positioned to integrate information about stimuli, actions, and rewards by tracking trial-by-trial outcomes of responses (Bryden, Johnson, Tobia, Kashtelyan, & Roesch, 2011; Hayden, Heilbronner, Pearson, & Platt, 2011; Heilbronner & Hayden, 2016). In our task, inhibition of the ACC rendered post-decision waiting times less sensitive to the strength of visual information and performance accuracy across trials, without affecting perceptual discrimination itself: i.e., impaired second order but left the first order processes intact (Miyamoto et al., 2017). Previous work in primates has demonstrated that confidence reports are informed by both decision difficulty and elapsed decision time (or reaction time; (Fetsch, Kiani, Newsome, & Shadlen, 2014; Kiani, Corthell, & Shadlen, 2014; Kiani & Shadlen, 2009)). Even in the absence of a change in decision accuracy, longer reaction times are associated with lower confidence. In the present work, we demonstrate that the same effect is present in rats and is also supported by the ACC. Finally, we found that ACC inhibition decreased metacognitive efficiency, or the trial-by-trial correspondence between decision accuracy and waiting times. In humans, a similar effect has been reported for perturbations of activity in the dorsolateral prefrontal cortex, which is shown to be important for visual metacognition (Rounis et al., 2010).

We note that waiting time is an indirect measure of confidence and as such, the effect of brain manipulations should be interpreted with caution. Firstly, several cortical and subcortical brain regions participate in reward timing (Bakhurin et al., 2017; Huertas, Hussain Shuler, & Shouval, 2015; Levy, Zold, Namboodiri, & Hussain Shuler, 2017; Murakami, Shteingart, Loewenstein, & Mainen, 2017). Secondly, an overall reduction in waiting time can result from an increased delay sensitivity or impulsivity and therefore may not be reflective of confidence *per se*. Here, we found that inhibition of the BLA renders rats less willing to wait overall. However, this effect of BLA inhibition was independent of the strength of visual evidence to make a perceptual decision. Furthermore, whereas inhibition of the ACC decreased metacognitive efficiency, inhibition of the BLA failed to change this measure. Thus, during perceptual decision making, the BLA may overall increase waiting time for reward, perhaps enabling other brain regions to interpret and/or act on ACC signals related to the strengths of visual information.

Our post-decision wagering paradigm mimics many features of foraging tasks that involve patch-leaving decisions. In rats, the ACC represents expected outcomes and signals errors in reward prediction, and is engaged when a change in the course of action is required and encodes information about rewards in remote locations (Bryden et al., 2011; Hyman, Holroyd, & Seamans, 2017; Mashhoori, Hashemnia, McNaughton, Euston, & Gruber, 2018). Similarly, in primates, the dorsal ACC participates in foraging decisions, signaling the value of leaving a patch in pursuit of other opportunities in the environment (Hayden et al., 2011). Furthermore, the dorsal ACC signals the value of the rejected option after the decision has been made (Blanchard & Hayden, 2014). Future research is needed to determine whether the impairment produced by ACC inhibition is specific to post-decision wagering tasks or will also manifest in opt-out tasks.

### Visual metacognition in rats

Recent work documents important similarities in visual information processing between rodents and primates, although species differences do exist (Meier & Reinagel, 2011, 2013; Reinagel, 2015). Pigmented rat strains, like the Long-Evans strain we studied here, have previously been used for vision research (Reinagel, 2015). Here, we found that rats also show high levels of visual metacognition, adjusting post-decision waiting times based on the uncertainty in perceptual decisions. This may allow direct comparison with the modality most often assessed in human and nonhuman primates while enabling easier, precise circuit manipulations.

### BLA and ACC in learning under perceptual uncertainty

We show that stimulus-response remapping is facilitated by perceptual certainty. Critically, *both* the BLA and ACC are required for faster learning when perceptual certainty is strong enough to improve learning. Considering that the BLA only shifts confidence readout, the observed reversals of all benefits of confidence on learning strategies and consistency in following a stimulus-response rule after BLA inhibition suggest a direct role of BLA in learning under uncertainty. That is, if the influence of BLA on learning was due to shifting confidence readout we would expect a bias in a certain direction and not reversal of all effects. In contrast, the effects of ACC seem to work through distorting confidence readout because its inhibition mainly attenuated the effect of confidence on learning.

Our results are also consistent with previous observations that the ACC-BLA circuit adjusts the levels of attention directed at environmental cues for learning based on prediction errors (Bryden et al., 2011). More specifically, it has been shown in rats that there is strong attention-related activity in the ACC during the entire trial following unexpected changes in reward and is most pronounced prior to and during outcome-predictive cues (Bryden et al., 2011). In contrast, unsigned reward prediction errors in the BLA may serve as attention signals, occurring at the time of unexpected reward delivery and omission (Roesch, Calu, Esber, & Schoenbaum, 2010). The ACC and BLA share direct and indirect bi-directional projections and the activity in this circuit appears to be required for adaptive learning under conditions of uncertainty in the visual cues guiding decisions or perhaps under more general cases of learning under uncertainty (Farashahi et al., 2017; Stolyarova & Izquierdo, 2017; Soltani & Izquierdo, 2019).

## Methods

### Subjects

In total 31 male outbred Long Evans rats (Charles River Laboratories, Crl:LE, Strain code: 006) were used in the experiments. The housing room in the vivarium was maintained under a reversed 12/12 h light/dark cycle at 22°C and all behavioral testing was conducted during rats’ active phase, of the dark portion of the cycle (between 08:00 and 18:00h). Rats remained undisturbed for 3 days after arrival to our facility to acclimate to the vivarium. Each rat was then handled for a minimum of 10 min once per day for 5 days. Following handling, rats underwent stereotaxic surgery to express inhibitory Designer Receptors Exclusively Activated by Designer Drugs (DREADDs; or control null virus to express only a fluorescent protein but no mutant receptors) and allowed to recover for three weeks. Rats were subsequently food-restricted to ensure motivation to work for food for one week prior to and during the behavioral testing, while water was available *ad libitum* except during behavioral testing. All rats were pair-housed at arrival and separated on the last day of handling to facilitate post-surgical recovery and minimize aggression during food restriction. We ensured that rats did not fall below 85% of their free-feeding body weight, and we saw a significant increase in rat body weight throughout the prolonged behavioral testing. On the last two days of food restriction prior to behavioral training, rats were fed 20 sugar pellets in their home cage to accustom them to the food rewards. For each experiment, rats were randomly assigned into groups, with the exception of assignment into high-confidence (HC) and low-confidence (LC) conditions for reversal learning as detailed below. All procedures were approved by the Chancellor’s Animal Research Committee at the University of California, Los Angeles.

### Viral constructs

We used inhibitory (Gi-coupled) DREADDs on a CaMKIIa promoter to transiently inactivate projection neurons in the ACC and BLA during performance on the behavioral task. An adeno-associated virus AAV8 driving the hM4Di-mCherry sequence under the CaMKIIa promoter was used to transduce ACC or BLA neurons with DREADDs (*AAV8-CaMKIIa-hM4D(Gi)-mCherry*, packaged by Addgene). A virus lacking the hM4Di DREADD gene (*AAV8-CaMKIIa-EGFP*, packaged by Addgene) was used as a null virus control. There were four experimental groups of rats: the active virus in BLA (n=8), the active virus in ACC (n=7), the null virus in BLA (n=8), and the null virus in ACC (n=8). The groups with the null virus expressed in brain regions of interest allowed us to control for virus exposure, non-specific effects of surgical procedures and subsequent injections on behavior.

### Surgery

All surgeries were performed using aseptic stereotaxic techniques under isoflurane gas anesthesia (5% in O_2_ during induction and 2–2.5% in O_2_ for maintenance). After being placed into a stereotaxic apparatus (David Kopf; model 306041), the scalp was incised and retracted. The skull was then leveled to ensure that bregma and lambda were in the same horizontal plane. Small burr holes were drilled in the skull to allow cannulae with an injection needle to be lowered into the BLA (the injection needle extended 1mm below the cannulae and its tip was at AP: −2.5; ML: ±5.0; DV: −7.8 (0.1μl) and −8.1 (0.2μl) from skull surface) or ACC (0.3 μl, AP = +3.7; ML= ±0.8; DV = −2.6). The injection needle was attached to polyethylene tubing connected to a Hamilton syringe controlled by a syringe pump. The viruses were infused bilaterally at a rate of 0.1 μl/min. For the BLA, the ventral infusion was administered first (at −8.1) followed by the dorsal site (−7.8) since our prior experiments demonstrated more precise targeting with this approach. There was no waiting time between the two infusions for BLA. After the last viral infusions in BLA or single infusion in ACC, the needle was left in place for 10 minutes to allow for diffusion of the virus, after which the cannulae were slowly lifted out of the brain and the wounds stapled. Each surgery took approximately 40 min. All rats were given a three-week recovery period prior to food restriction and subsequent behavioral training. Carprofen (5mg/kg, s.c.) was administered for 5 days postoperatively to minimize pain and discomfort. Behavioral measures of discomfort and conditions of the wounds were monitored daily, and all surgical staples were removed within 7-10 days after surgeries depending on a rat’s recovery.

### Electrophysiological confirmation of DREADDs

Separate rats were prepared with ACC DREADDs using identical surgical procedures to the main experiments. Slice recordings did not begin until at least three weeks following surgery to allow sufficient hM receptor expression. Slice recording methods were similar to those previously published (Babiec, Jami, Guglietta, Chen, & O’Dell, 2017). Three rats were deeply anesthetized with isoflurane and decapitated. The brain was rapidly removed and submerged in ice-cold, oxygenated (95% O_2_/5% CO_2_) artificial cerebrospinal fluid (ACSF) containing (in mM) as follows: 124 NaCl, 4 KCl, 25 NaHCO_3_, 1 NaH_2_PO_4_, 2 CaCl_2_, 1.2 MgSO_4_, and 10 glucose (Sigma-Aldrich). 400-μm-thick slices containing the ACC were then cut using a Campden 7000SMZ-2 vibratome. Slices from the site of viral infusion were used for inhibitory G-protein (G)_i_ validation. Expression of mCherry was confirmed post-hoc. Slices were maintained (at 30°C) in interface-type chambers that were continuously perfused (2–3 ml/min) with ACSF and allowed to recover for at least 2 hours before recordings. Following recovery, slices were perfused in a submerged slice recording chamber (2–3 ml/min) with ACSF containing 100 μM picrotoxin to block GABA_A_ receptor-mediated inhibitory synaptic currents. A glass microelectrode filled with ACSF (resistance = 5– 10 MΩ) was placed in layer 2/3 ACC to record field excitatory postsynaptic synaptic potentials and population spikes elicited by layer 1 stimulation delivered using a bipolar, nichrome-wire stimulating electrode placed near the medial wall in ACC. Stimulation intensity (0.2 msec duration pulses delivered at 0.33 Hz) was set to the minimum level required to induce reliable population spiking in ACC. Once reliable responses (measured as the area of postsynaptic responses over a 4 second interval) were detected, baseline measures were taken for at least 10 minutes, followed by a 20 minutes bath application of 10 μM CNO. Unless noted otherwise, all chemicals were obtained from Sigma-Aldrich.

### Behavioral training

Behavioral training was conducted in operant conditioning chambers (Model 80604, Lafayette Instrument Co., Lafayette, IN) that were housed within the sound- and light- attenuating cubicles. Each chamber was equipped with a house light, tone generator, video camera, and LCD touchscreen opposing the pellet dispenser. The pellet dispenser delivered 45-mg dustless precision sucrose pellets. Software (ABET II TOUCH; Lafayette Instrument Co., Model 89505) controlled the hardware. All testing schedules were customized in ABET by our group and can be requested from the corresponding author. During habituation, rats were required to eat five pellets out of the pellet tray inside of the chambers within 15 min before exposure to any stimuli on the touchscreen. They were then progressively trained to respond to visual stimuli presented on the screen, to initiate the trial, report the orientation of the visual stimulus (vertical or horizontal) by nosepoking left or right on a white square stimulus, and wait for rewards.

### Behavioral Testing and experimental paradigm

A rat first initiated each trial by nosepoking a bright white square in the center of the screen. The initiation stimulus then disappeared, and a rat was briefly (1s) presented with a vertical (V) or horizontal (H) Gabor patch embedded in noise, and required to report the orientation (H or V) based on a complementary stimulus-response rule, e.g., H→left and V→right. These spatial responses were made by nosepoking the right or left compartments of the touchscreen that became illuminated after the disappearance of the oriented visual stimulus. We altered two properties of the visual stimuli to manipulate their ambiguity. First, we changed the signal-to-noise ratio (SNR), defined as the ratio of the contrast of the Gabor patch relative to the contrast of the added Gaussian noise. Second, we changed the overall contrast of both the Gabor patch and the added noise for a given SNR. Gratings were 200 pixels square, with spatial frequency 20 px/cycle. For training, gratings were presented at 100% contrast. For testing, gratings were embedded in white noise as follows. To create different contrasts designed to produce a range of performance (measured by *d*′) and confidence (measured by waiting time) responses such that HC and LC conditions could be established, animals performed the task on 40%, 60%, and 80% maximum contrast Gabor patches embedded in noise also with three possible levels of increasing contrast, for nine possible full-factorial combinations in total. This method of constant stimuli (Macmillan & Creelman, 2004) facilitated selection of a pair of stimuli from these nine levels such that the animal had produced matched perceptual performance capacity (*d*′) but different waiting time in HC and LC conditions.

Correct choices were reinforced probabilistically after a randomly assigned delay: 70% of correct responses resulted in reward delivery. Time to reward delivery was drawn from an exponential distribution with mean of 8 sec (see **Supplementary Figure 4** for an example and the average distributions of reward delivery) and on trials with no reward, the trial ends after 40 sec of no re-initiation occurs. Specifically, following stimulus discrimination, rats expressed their confidence by time wagering: they could wait a self-timed delay in anticipation of reward or initiate a new trial similar to previous work by the Kepecs lab (Lak et al., 2014). The initiation stimulus appeared on the touchscreen 2 sec after a rat indicated its choice. This delay was imposed to prevent non-discriminant responding. We define the time that the animal waited before re-initiating a trial as the waiting time (see **Supplementary Figure 4** for an example and the average the distributions of waiting time).

Following fully learning the task and testing on the perceptual decision-making with re-initiation (confidence report), rats were randomly assigned to a high confidence (HC)- or low confidence (LC) condition and experienced a reversal in the stimulus-response rule. In order to determine the visual stimuli for HC and LC conditions for each rat, we selected two SNR and contrast levels that had equal discrimination accuracy (*d*′) and reinforcement history and were different only in confidence levels measured by waiting time. After determining discrimination-matched stimuli for each rat, rats were randomly assigned to LC and HC conditions and the corresponding stimulus was used for each rat based on the assigned condition. After the reversal in stimulus-response rule, rats were no longer offered an option to re-initiate the trial, but were required to wait a random delay before reward delivery or the end of the trial (on no-reward trials) following a response. This was to simplify the re-learning and ensure rats were not adopting a complex strategy due to the availability of the re-initiation option.

To study the contributions of BLA and ACC to decision-making and learning under perceptual uncertainty, we used a within-subject design: rats were given vehicle injections, CNO injections, and no injection (prior to reversal). The order of CNO and vehicle injections was counterbalanced. Therefore, a subset of behavioral sessions were preceded by inactivation of ACC or BLA pyramidal neurons via peripheral (3mg/kg; i.p.) administration of clozapine-n-oxide (CNO) 10 min prior to the testing. The injections were administered in rats’ housing room. Due to the long duration of pretraining on our task, all CNO injections were administered at least 12 weeks following the surgery, ensuring sufficient virus transduction and receptor expression. On another subset of sessions, rats received vehicle to control for behavioral effects of the stress of injections. All rats received 2-day wash-out period between drug conditions and the order of injections was counterbalanced across rats.

### Histology

Rats were euthanized within 90 min following the last testing session with an overdose of Euthasol (Euthasol, 0.8 mL, 390 mg/mL pentobarbital, 50 mg/mL phenytoin; Virbac, Fort Worth, TX), were transcardially-perfused, and their brains removed for histological processing. Brains were fixed in 10% buffered formalin acetate for 24 hours followed by 30% sucrose for 5 d. To visualize hM4Di-mCherry and -EGFP expression in BLA or ACC cell bodies, free-floating coronal sections were mounted onto slides and coverslipped with mounting medium for DAPI. Slices were visualized using a BZ-X710 microscope (Keyence, Itasca, IL), and analyzed with BZ-X Viewer and analysis software.

### Signal detection theory analyses

According to standard signal detection theory, *d*′ measures how well a subject’s perceptual decisions track physical stimuli. *d*′ is preferred to other discrimination accuracy measures (i.e. percent or probability correct) because it accounts for biases such as side bias, stimulus preference (Odegaard et al., 2018). Extending the same approach to confidence measures, metacognitive sensitivity measures how well confidence tracks the likelihood that a perceptual decision is correct, and like *d*′, can also be formulated to account for bias. Specifically, Maniscalco & Lau (2012) have proposed meta-*d*′ to measure metacognitive sensitivity on the same scale as *d*′ so that one can calculate the ratio between the two (meta-*d*′/*d*′) to assess the metacognitive efficiency of a subject. They defined the task of classifying stimuli as a Type 1 task, whereas the rating of confidence in this classification as a Type 2 task. Meta-*d*′/*d*′ varies between 0 and 1, where 0 indicates that the rat’s trial-by-trial waiting times (i.e. confidence) do not correspond with trial accuracy and 1 indicates that the rat’s Type 2 capacity is exactly matching its Type 1 sensitivity. In other words, subjects could wait a longer time before re-initiating a trial when their response is correct, and wait less time before re-initiating when the response is incorrect, considering the limitation in discrimination. Meta-*d*′/*d*′, therefore, should be larger than 0 and also significantly less than 1; a meta-*d*′/*d*′ of 1 would indicate ideal or optimal metacognitive behavior. Compared to commonly used Type 2 receiver operating characteristic (ROC) analysis, the meta-*d*′/*d*′ approach has the advantage of allowing one to isolate the effects of confidence on behavior from basic perceptual performance capacity. To calculate meta-*d*′, we used MATLAB (MathWorks, Natick, MA) functions freely available at http://www.columbia.edu/~bsm2105/type2sdt/.

### Data analyses

We used MATLAB (MathWorks, Natick, MA; Version R2018b) for data and statistical analyses. For trial-by-trial learning analyses, we included data from all 15 rats that completed a total of 127,303 trials in 603 sessions. Learning occurred in a mixed design with three within-subject/repeated-measures of stage (vehicle, inhibition, no-injection) and three between-subjects-(group) conditions of vehicle, ACC inhibition, and BLA inhibition. All 15 rats experienced the vehicle and no-injection conditions. However, 8 and 7 rats experienced BLA and ACC inhibition conditions, respectively. Since it was possible that a rat received reward prior to “intended” re-initiation of a trial, we excluded rewarded trials in the analysis of waiting time and reaction time. Furthermore, we excluded the trials in which reaction time deviated from the mean of the reaction time of the session by more than three times the standard deviation. This criterion resulted in removal of 1.1% of the trials.

Unlike the analyses presented in the preceding sections with n=7 or 8 for each targeted brain region, statistical results for after reversal were restricted to 3 or 4 rats per group due to the additional HC and LC conditions (n=4 rats in each of the HC and LC conditions with the BLA as the targeted region, n=4 rats in the HC condition and n=3 rats in the LC condition with the ACC as the targeted region). This constraint was a consequence of the experimental (within-subject) design and longitudinal nature of pre-training on the task, followed by extensive learning. Not including pretraining, the mean number of sessions to complete both the initial and re-learning part of the experiments was 40–210 trials on average, in each session.

For comparisons of learning strategies and consistency following in following a stimulus-response rule across different experimental conditions, we used permutation test. More specifically, we first calculated the actual probability of using a strategy (e.g., Win-Stay) in the observed data and then permuted this data 10000 times to construct the permutation or null distribution. We then calculated the probability of obtaining the observed value for use of the strategy based on the null distribution from which we estimated p-values (Hesterberg et al., 2005).

To compare the rate of learning after reversal we used an exponential function to fit the learning curve from all rats in a given experimental condition:

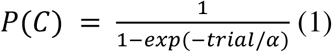

where *α* is the learning parameter.

Finally, we showed the only significant difference between the HC and LC conditions prior to reversal learning was waiting time for the assigned contrast-SNR pairs that were administered after reversals. Since contrast is correlated with SNR, we used a stepwise regression to find the variables which contributed significantly in explaining the response variables (waiting time, reward history). Confidence condition was entered as a predictor variable in a GLM along with other variables (SNR, contrast, targeted brain region, and trial accuracy) in a stepwise manner to observe which of these increased adjusted R-squared significantly for the response variables.

### Rule-based Repetition index (RRI)

In order to examine the consistency in following a stimulus-response rule on two consecutive trials, we used a repetition index that was previously introduced to capture tendency to repeat the same choice beyond what is expected by chance (Soltani et al., 2013), and extended it to selection based on response rules. Specifically, we computed the probability that the same rule (either correct or incorrect) was used on two consecutive trials, *p*(StayRule), and subtracted the tendency to repeat the same rule on two consecutive trials due solely to chance to arrive at the rule-based repetition index:

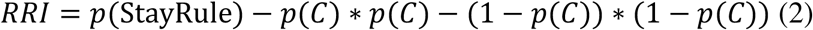

where *p*(C) is the probability of choosing the correct rule. Therefore, unlike other perseveration indices (Izquierdo et al., 2006), the RRI accounts for the probability of using the same rule on consecutive trials by chance.

## Acknowledgements

We thank Dr. Brian Maniscalco for helpful comments on these data. This work was supported by CRCNS R01DA047870 (Soltani and Izquierdo), UCLA’s Division of Life Sciences Recruitment and Retention Fund, and UCLA Academic Senate Grant (Izquierdo). We acknowledge the Charles E. and Sue K. Young Fellowship (Stolyarova). We also thank the Staglin Center for Brain and Behavioral Health for enabling the use of equipment for microscopy imaging.

## Supplementary analyses

### Trial re-initiation is only affected by SNR

In order to ensure that our drug manipulations did not change the tendency of the animals to re-initiate a new trial, we used a GLM to examine whether the number of re-initiations was affected by either vehicle or CNO administration, or by either the BLA and ACC as the targeted brain region, or SNR of the stimuli. Importantly, we found that only SNR significantly affected the number of trial re-initiations (GLM; *F*(7,871) = 13.5, *p* = 1.14 × 10^−16^, adjusted *R*^2^ = 0.0908; ratio: *β* = −2.26, *p* = 0.00026), indicating that re-initiation mainly depended on the strength of the visual information and thus perceptual uncertainty.

### Relationship between waiting time and reaction time in different experimental conditions

The confidence intervals (95%) for the slopes of linear regressors indicated that the negative correlations between waiting time and reaction time in the two control conditions (vehicle and no-injection prior to reversal) were not significantly different, but were significantly different following CNO administration. More specifically, the confidence intervals for the slopes are as follows: vehicle administration, [−2.7061, −2.3402]; ACC inhibition, [−0.8850, −0.3630]; BLA inhibition, [−1.6288, −1.2418]; no-injection, [−2.7086, −2.3478] (**Supplementary Figure 7C,D**). Thus, the correlation between these measures was still negative but weaker in both ACC and BLA inhibition conditions.

### Vehicle administration does not change the behavior

To show that vehicle administration alone does not produce the observed behavioral changes, we compared the effect of vehicle administration and no-injection on waiting time and reaction time. Similar to vehicle administration, we found that in the sessions in which no injection was administered, waiting times were longer on trials with a larger SNR (GLM; no-injection: *F*(7,904) = 212, *p* = 1.26 × 10^−185^, adjusted *R*^2^ = 0.618; *r* = 1.75, *p* = 2.46 × 10^−15^; **Supplementary Figure 6**). Furthermore, waiting times were negatively correlated with reaction times on a trial-by-trial basis (GLM; ACC inhibition: slope= −0.62, *p* = 2.83 × 10^−6^; BLA inhibition: slope= −1.43, *p* = 2.88 × 10^−47^; no-injection: slope= −2.53, *p* = 3.02 × 10^−162^).

### Absence of non-specific effects of virus exposure

In the main body of the manuscript we presented data on behavior following vehicle and CNO administration in rats with DREADDs expressed in the BLA and ACC. Here, we include data demonstrating that these impairments are not due to non-specific effects of surgery/virus exposure. Therefore, in this section we compare null (EGFP) and active (DREADDs) virus following vehicle administration on the major measures of the task such as waiting time, reaction time, performance, *d*′, and meta-*d*′.

We first observed that the type of virus (null vs. active) had no significant effect or interaction with trial type (correct vs. incorrect) and/or SNR values (GLM; *F*(7,178) = 95.7, *p* = 6.12 × 10^−57^, adjusted *R*^2^ = 0.782; virus type: *β* = 1.61, *p* = 0.3; virus type × trial type: *β* = −0.16, *p* = 0.94; virus type × ratio: *β* = −0.008, *p* = 0.98; virus type × trial type × ratio: *β* = −0.35, *p* = 0.62). Similarly, we did not find a significant effect of virus type in either main effect or interactions with trial type and SNR (GLM; *F*(7,178) = 177, *p* = 9.02 × 10^−77^, adjusted *R*^2^ = 0.87; virus type: *β* = 0.09, *p* = 0.23; virus type × trial type: *β* = −0.13, *p* = 0.25; virus type × ratio: *β* = −0.03, *p* = 0.15; virus type × trial type × ratio: *β* = 0.03, *p* = 0.44).

Next, we analyzed the effect of virus type on performance measures of probability of correct response, *d*′, and meta-*d*′. We observed no significant effect of virus type or interaction with SNR on probability of correct response (GLM; *F*(3,89) = 70.3, *p* = 2.16 × 10^−23^, adjusted *R*^2^ = 0.693; virus type: *β* = −0.03, *p* = 0.15; virus type × ratio: *β* = 0.01, *p* = 0.16). Furthermore, we observed that the type of virus had no significant effect or interaction with SNR values on *d*′ (GLM; *F*(3,89) = 4.85, *p* = 0.003, adjusted *R*^2^ = 0.112; virus type: *β* = −0.13, *p* = 0.64; virus type × ratio: *β* = −0.044, *p* = 0.63). Finally, we observed no significant effect of virus type or interaction with ratio on meta-*d*′ (GLM; *F*(3,89) = 1.18, *p* = 0.32, adjusted *R*^2^ = 0.006; virus type: *β* = −0.25, *p* = 0.53; virus type × ratio: *β* = 0.02, *p* = 0.88). These results show that the observed impairments through CNO administration was specific to the active (DREADDs) virus we used in the experiment.

**Figure S1.**
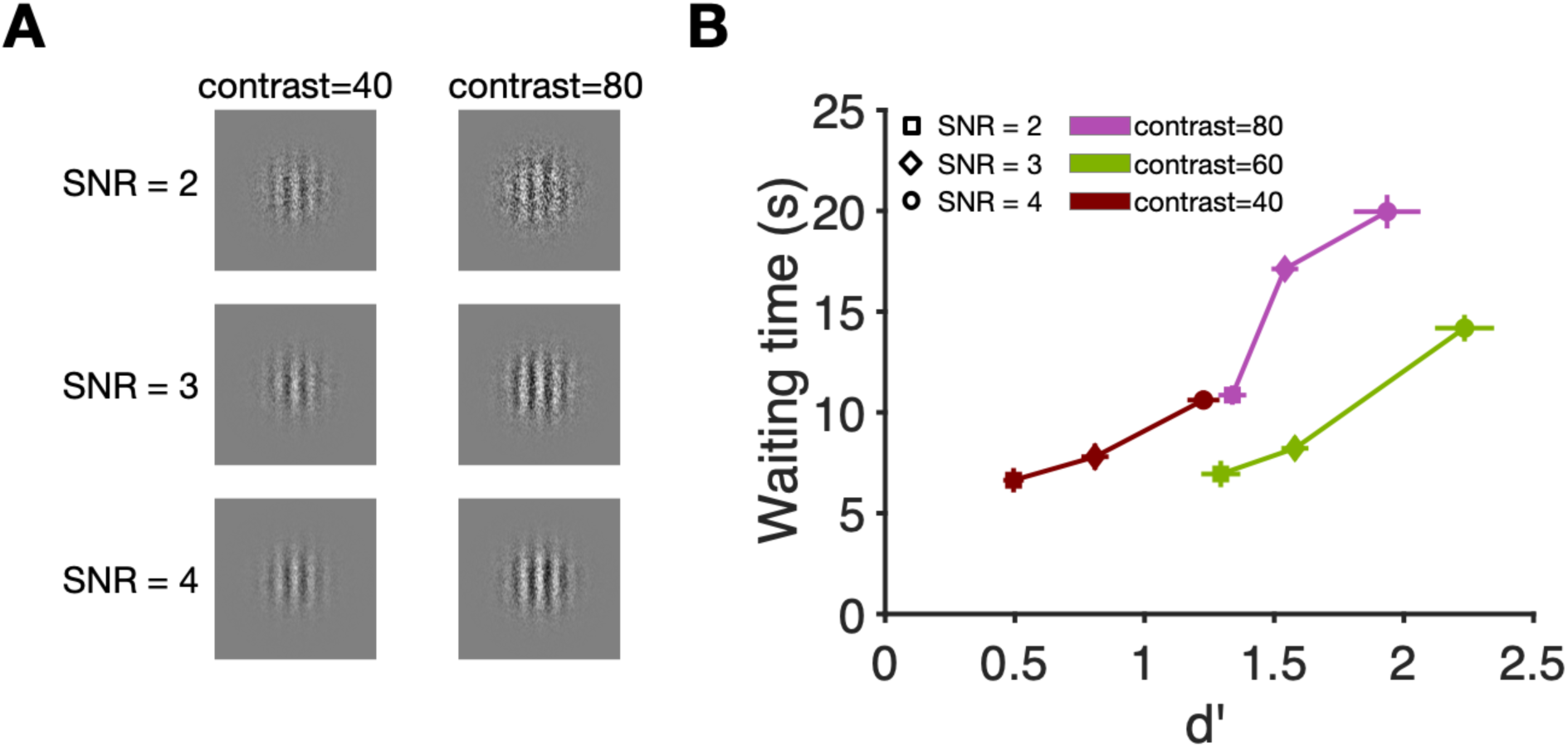
SNR, contrast of the visual stimuli, and waiting time and discrimination performance as a function of those manipulations. **(A)** Examples of visual stimuli with more (80) or less (40) contrast with different signal-to-noise (SNR) ratio, reflecting the strength of the visual signal (4, most discriminable; 3, moderately discriminable; 2, least discriminable). (**B**) Waiting time and *d*′ increases with SNR for any value of contrast.

**Figure S2.**
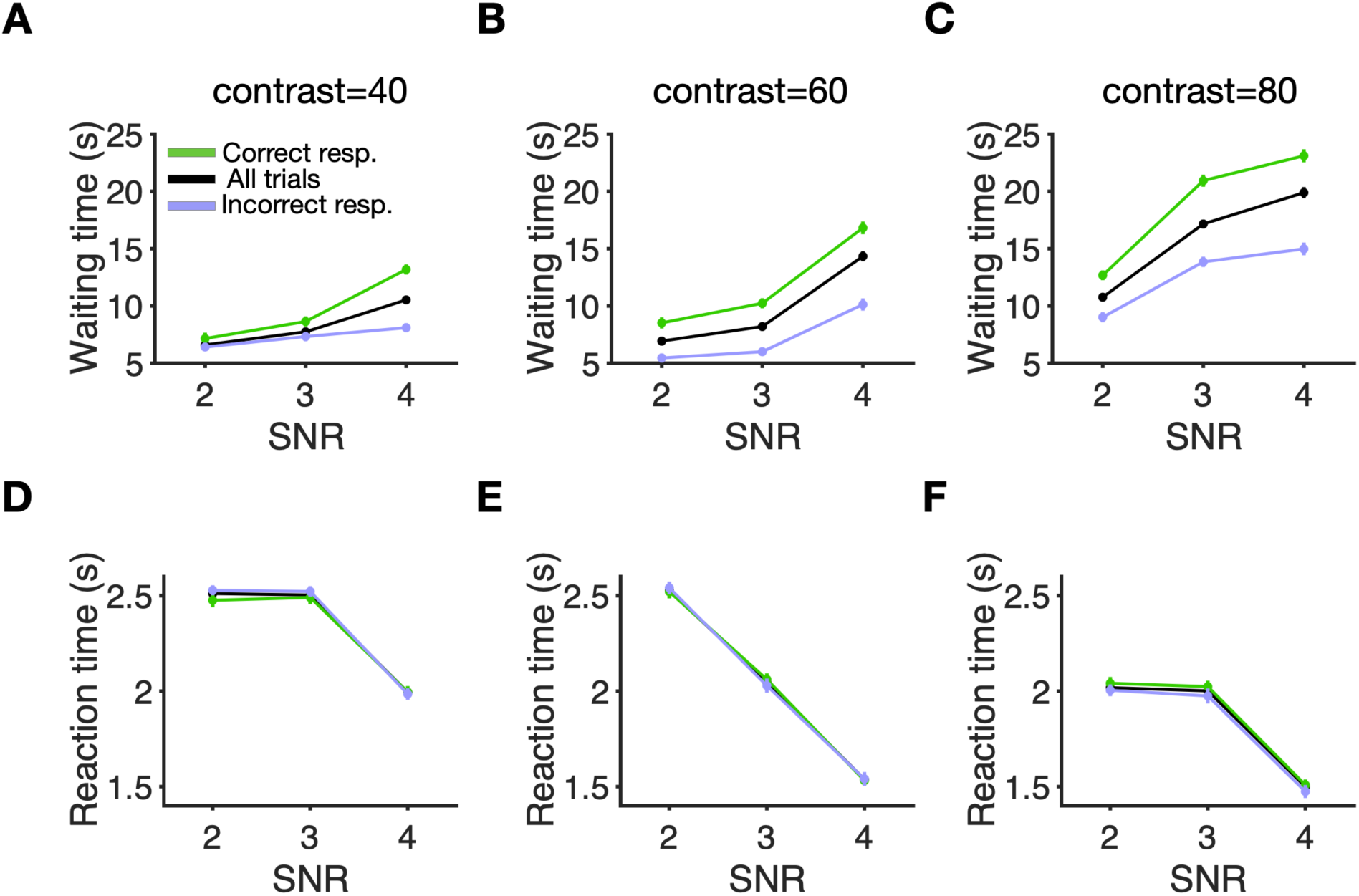
Waiting time increases with SNR for any value of contrast whereas reaction time dependence on SNR was strongly modulated by contrast. Plotted are the waiting time and reaction time as a function of SNR for different values of contrasts, and separately for correct and incorrect trials. **(A-C)** Plotted is the waiting time for all trials as a function of three contrast levels (40, weaker to 80, stronger) and different signal-to-noise (SNR) ratio, reflecting the strength of the visual signal (4, most discriminable; 3, moderately discriminable; 2, least discriminable) for correct responses (green), incorrect responses (blue), and all trials (black). (**D-E**) Same as in panel A-C but for reaction times. Error bars show the S.E.M. over sessions (typically smaller than the symbols).

**Figure S3.**
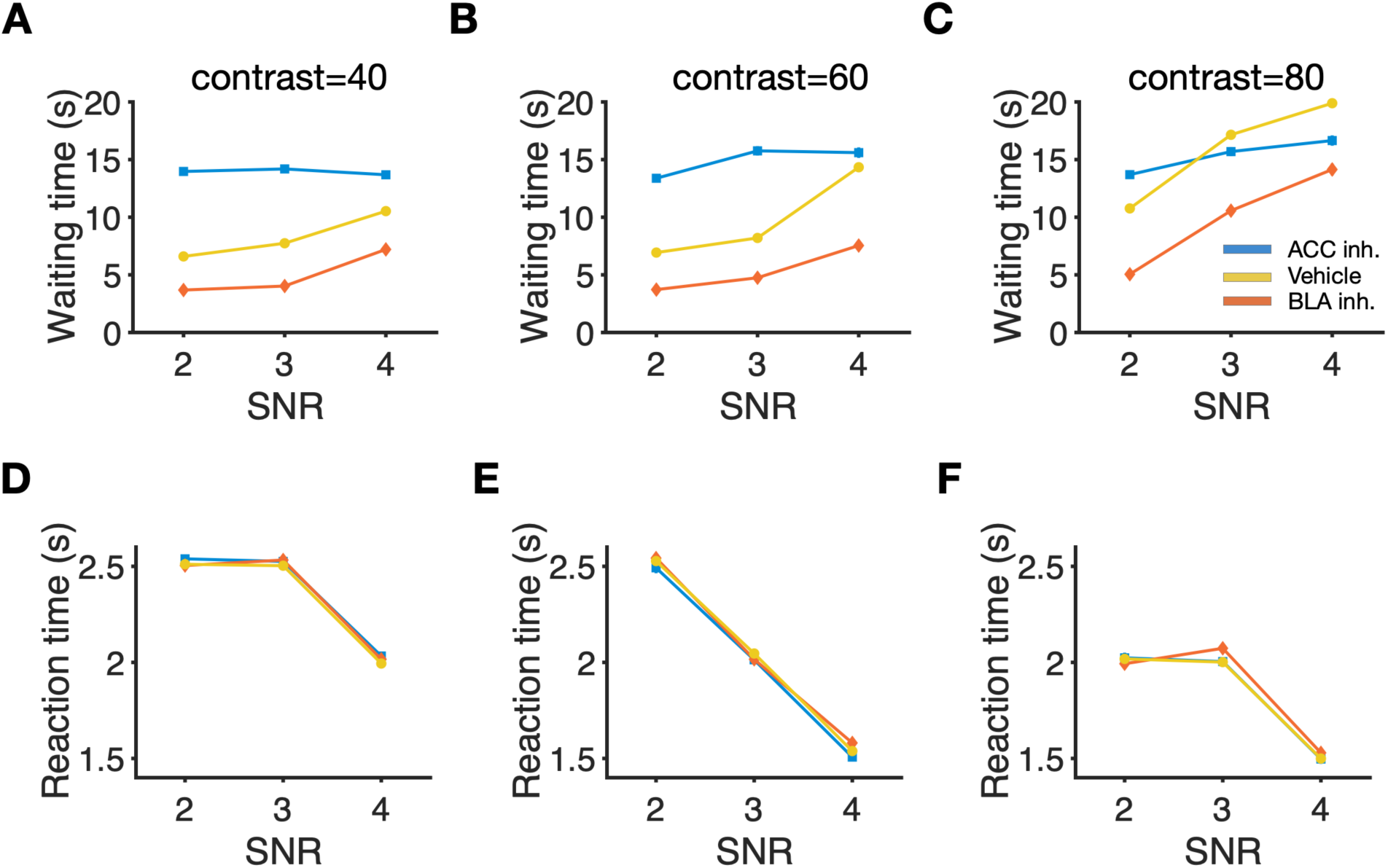
ACC and BLA inhibition influenced waiting time for all values of contrast whereas reaction time was not sensitive to these manipulations. Plotted are waiting time and reaction time as functions of SNR for different values of contrast after the inhibition of ACC and BLA. Conventions are the same as in Figure 3. Overall, ACC inhibition rendered waiting time insensitive to SNR for all contrast values such that at high contrast, waiting time in ACC-inhibited rats fell below the control condition. In contrast, BLA inhibition, reduced waiting time for all contrast values without reducing the sensitivity to SNR.

**Figure S4.**
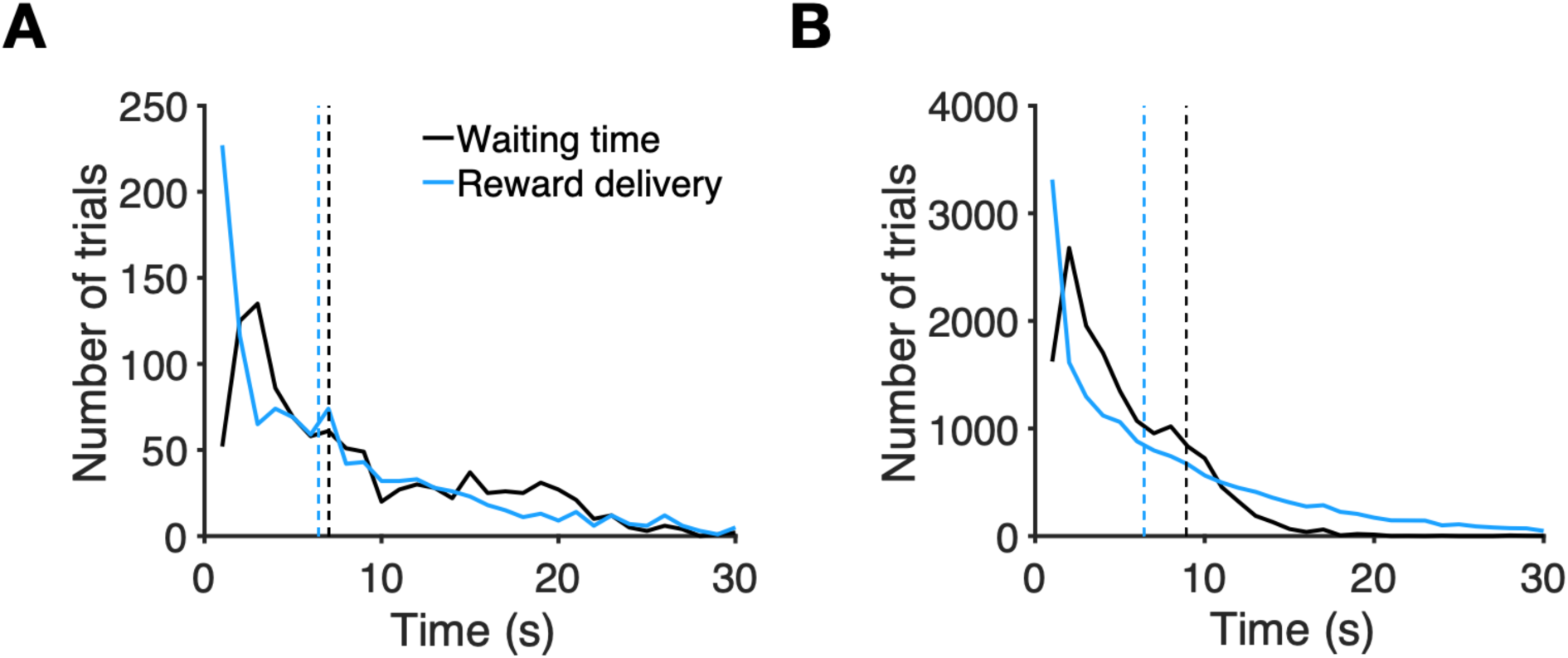
Rats’ waiting time tracked distributions of the statistics of reward delivery time. Plotted are the distributions of reward delivery time (blue) and waiting time (black) in an example individual rat (A) and across all rats (B). The dashed lines show the median of the reward delivery time distributions and of waiting time, respectively. The waiting time distribution generally followed that of the reward delivery time.

**Figure S5.**
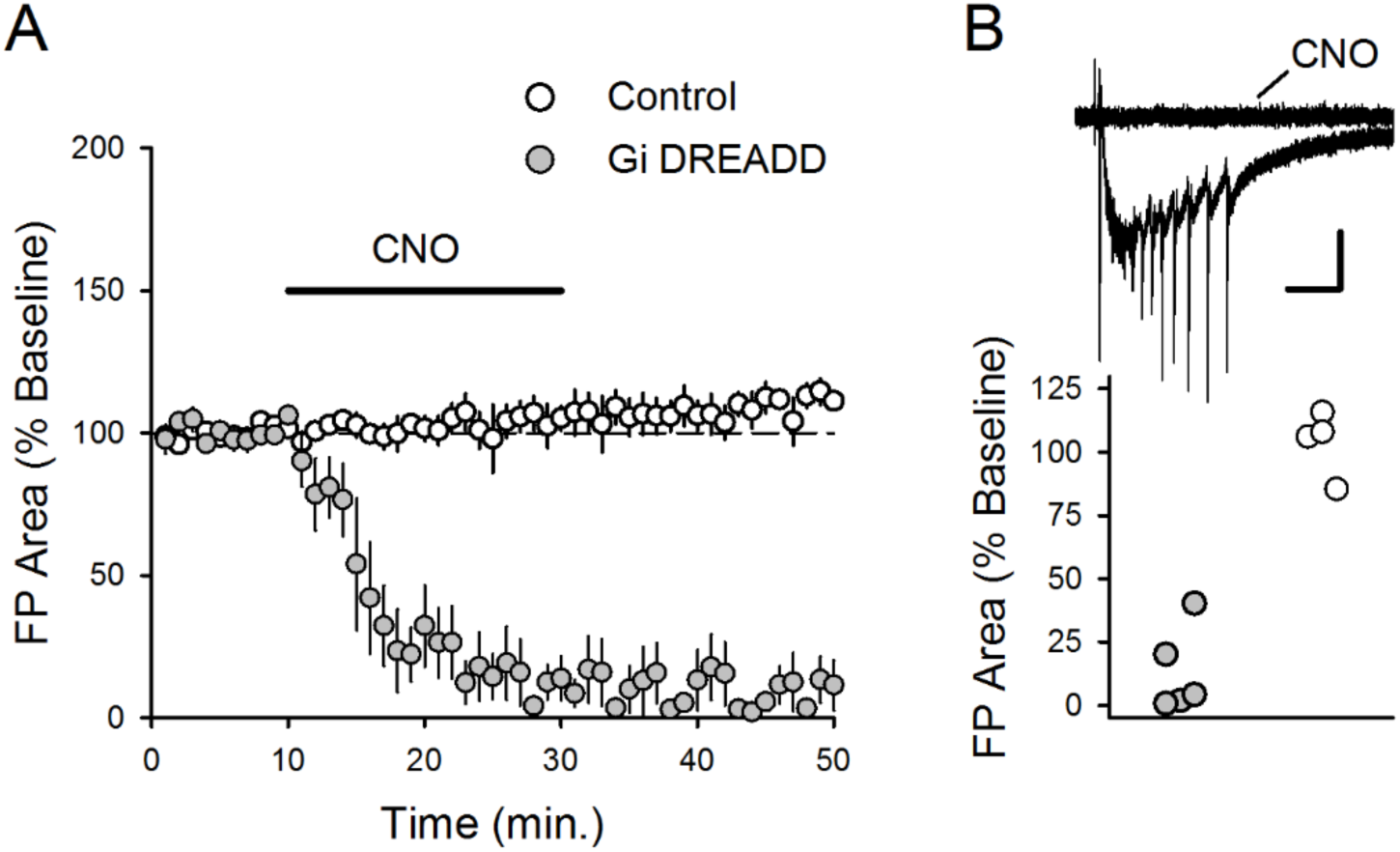
Electrophysiological effect of bath application of CNO in slice. (**A**) Inhibition of field potentials (FP) elicited by layer I stimulation followed by bath application of CNO (10 μM, indicated by the bar) in a control and DREADD preparation. Plot shows the FP area (normalized to baseline) over time. CNO was applied at *t* = 10 min (**B**) Top: Traces show superimposed responses elicited before and after CNO application in transfected slices. Calibration bars: 0.5 mV, 500 milliseconds. Bottom: Points show FP area (normalized to baseline) at end of CNO application (averaged over last 5 min) from individual slices. Application of CNO strongly suppressed FPs in transfected slices (filled circles; FP area was reduced to 13.4 ± 8% of baseline; paired t-test for comparison to baseline, *t*(4) = 11.333, *p* = 3.46×10^−3^, *n* = 5 slices from 3 rats) but had no effect on responses in non-transfected slices (open circles; FP area was 103.7 ± 7% of baseline; paired t-test for comparison to baseline, *t*(3) = 0.578, *p* = 0.604, *n* = 4 slices from 3 rats).

**Figure S6.**
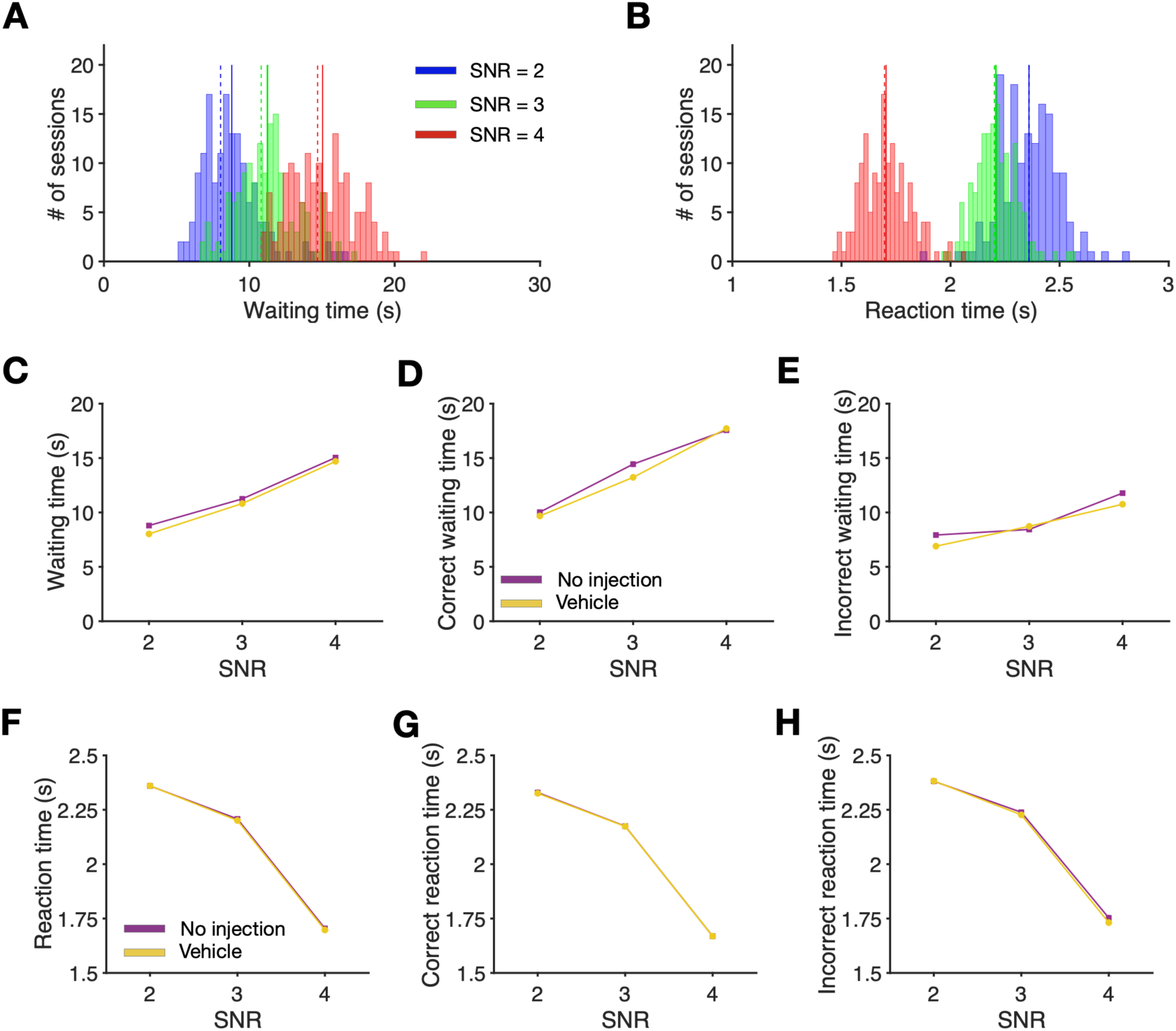
Behavior is similar following vehicle administration and no-injection control. **(A)** Waiting time in sessions with no injection increases similarly to sessions following vehicle administration. Plotted are the distributions of the waiting time for each SNR:2 (blue), 3 (green), and 4 (red), following no injection. Solid lines show median of each distribution. Dashed lines show the median of the same distributions but following vehicle administration as is shown in Figure 2A. **(B)** Same as in panel A but for reaction times. **(C-E)** Following no injection waiting times are similarly sensitive to the strength of sensory signal compared to sessions after vehicle administration. (C) Plotted is the waiting time for all trials as a function of SNR following vehicle administration (yellow), and no injection (purple). (D) Same as panel C but only on trials in which the correct choice was made. (E) Same as panel C but only on trials in which the incorrect choice was made. **(F-H)** Reaction time changes similarly following no injection compared to the sessions following vehicle administration. (F) Plotted is the reaction time for all trials as a function of SNR following vehicle administration (yellow), and no injection (purple). (G) Same as panel F but only on trials in which the correct choice was made. (H) Same as panel F but only on trials in which the incorrect choice was made. Error bars show the S.E.M. over sessions and they are usually smaller than the marker.

**Figure S7.**
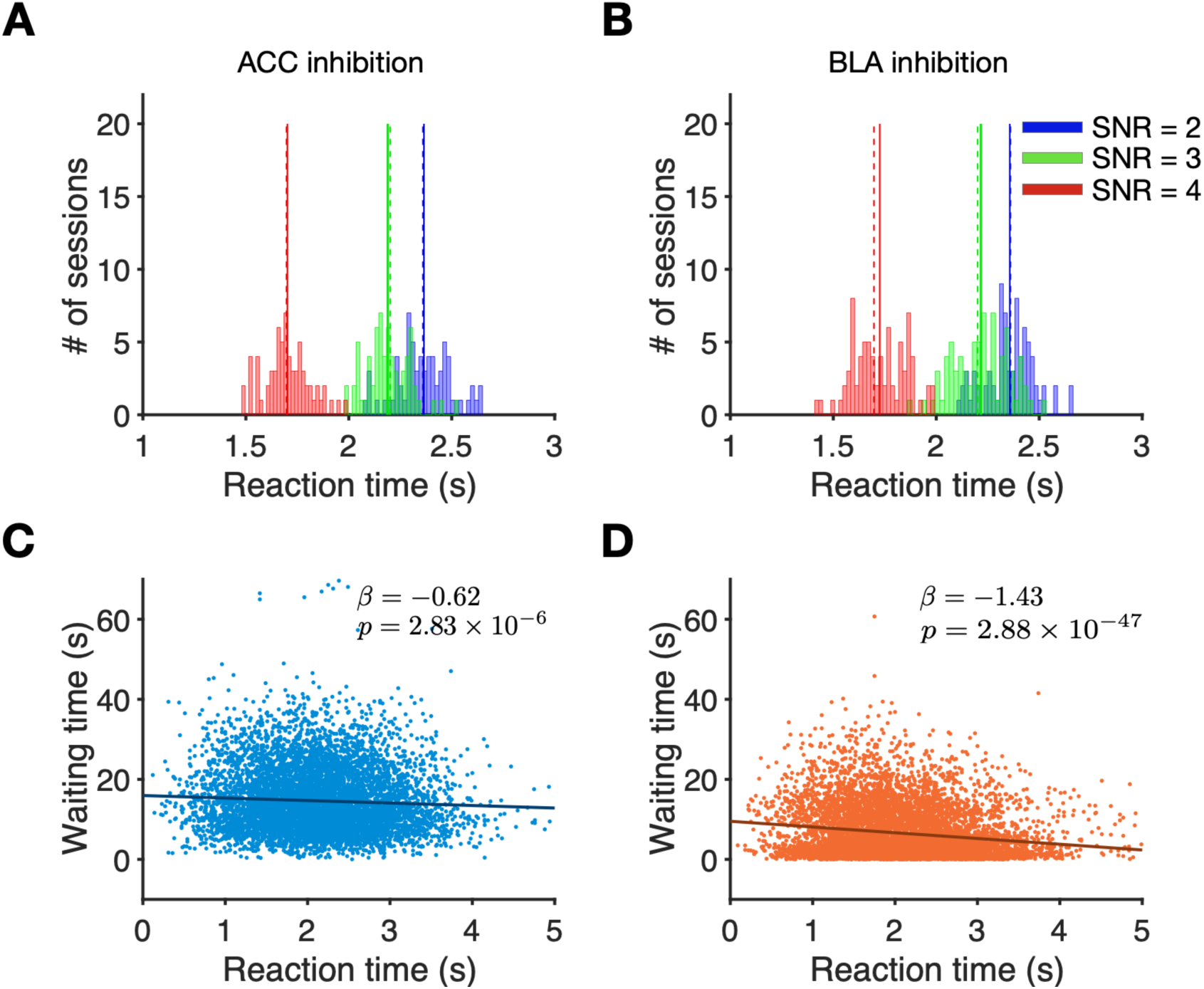
Reaction time is insensitive to inhibition of either ACC or BLA. **(A)** Reaction time does not change following ACC inhibition. Plotted are the distributions of the reaction time for each SNR:2 (blue), 3 (green), and 4 (red), following CNO administration of ACC. Solid lines show median of each distribution. Dashed lines show the median of the same distributions but following vehicle administration as is shown in Figure 2A. **(B)** Same as in panel A but for sessions following inhibition of BLA, via CNO injection. Solid lines show median of each distribution. Dashed lines show the median of the same distributions but following vehicle administration as is shown in Figure 2A. **(C)** Waiting time before re-initiation of a new trial is negatively correlated with reaction time to make a choice following inhibition of ACC. Waiting time is plotted as a function of the reaction time for all trials and all rats. Each data point is a trial in a session following vehicle administration. **(D)** Same as in panel C but for sessions following inhibition of BLA, via CNO injection.

